# Ascending inputs to inferior colliculus subdivisions reveal pathway-specific hybrid organization in the external cortex

**DOI:** 10.64898/2026.05.09.724030

**Authors:** Hailey C. Acosta, Michellee M. Garcia, Melanie Andrade-Muñoz, Michael R. Kasten, Paul B. Manis, Hiroyuki K. Kato

## Abstract

The inferior colliculus (IC) is a major hub of the auditory system, integrating ascending inputs from auditory and non-auditory brainstem nuclei before relaying information to the thalamus, and ultimately, the cortex. The IC is classically divided into the central nucleus (CIC) and surrounding shell regions, including the external (ECIC) and dorsal (DCIC) cortices. Although numerous tracing studies have described ascending inputs to the IC, most have either treated the IC as a uniform structure or distinguished only core and shell regions, leaving potential heterogeneity within the shell unresolved. Here, we performed retrograde tracing from distinct subdivisions of the mouse IC to compare their ascending input organization. By systematically targeting the CIC, DCIC, and both rostral and caudal ECIC and registering labeled neurons to a standardized brain atlas, we quantified input neurons across auditory and non-auditory brainstem structures. The CIC exhibited input patterns largely consistent with canonical pathways. In contrast, the DCIC received greater input from paralemniscal, periolivary, and non-auditory nuclei. Notably, the ECIC did not simply receive DCIC-like inputs or occupy an intermediate position between CIC and DCIC. Instead, it displayed a pathway-specific hybrid organization: inputs from the nuclei of the lateral lemniscus and superior olivary complex were largely CIC-like, whereas inputs from the cochlear nucleus and non-auditory regions resembled those of the DCIC. Furthermore, rostral and caudal ECIC showed overlapping but distinct input patterns. Together, these results reveal heterogeneous ascending connectivity across IC subdivisions and identify the ECIC as a hybrid interface that bridges CIC-like and DCIC-like ascending pathways.

**Significance Statement:** The inferior colliculus is often divided into a lemniscal core and nonlemniscal shell, but whether shell subdivisions share a common ascending input organization has not been directly tested. By systematically mapping ascending inputs to the central nucleus, dorsal cortex, and rostral and caudal external cortex, we show that the shell regions are not uniform. Instead, the external cortex combines core-like inputs from the nuclei of the lateral lemniscus and the superior olivary complex with dorsal cortex-like inputs from the cochlear nucleus and non-auditory regions. These findings revise the classical core-versus-shell framework and provide an anatomical basis for parallel midbrain pathways, in which the external cortex is positioned for fast auditory signaling while simultaneously receiving contextual and multisensory modulation.

## Introduction

The inferior colliculus (IC) is the principal midbrain hub of the auditory system that integrates ascending inputs from multiple brainstem nuclei before relaying information to the thalamus and cortex. Anatomically, the IC is commonly subdivided into the central nucleus (CIC), external cortex (ECIC), and dorsal cortex (DCIC), which differ in cytoarchitecture, connectivity, and sound response properties (Morest and Oliver, 1984; Faye-Lund and Osen, 1985; Oliver, 2005; Loftus et al., 2008). The CIC is generally regarded as the lemniscal or “core” subdivision, characterized by tonotopic organization, short-latency sound responses, and dense ascending inputs from major auditory brainstem nuclei, including the lateral lemniscus (LL), superior olivary complex (SOC), and cochlear nucleus (CN) (Beyerl, 1978; Roth et al., 1978; Adams, 1979; Brunso-Bechtold et al., 1981; Kelly et al., 1998). In contrast, the ECIC and DCIC have classically been grouped as “shell” or nonlemniscal subdivisions and are generally considered secondary regions with less prominent tonotopy, multimodal sensory responses, and stronger corticofugal modulation (Aitkin et al., 1978; Winer et al., 1998; Syka et al., 2000; Jain and Shore, 2006; Bajo et al., 2010; Wong and Borst, 2019; Blackwell et al., 2020; Quass et al., 2024; Ibrahim et al., 2025). Although these functional distinctions suggest underlying differences in ascending connectivity, these inputs have not been systematically compared across IC subdivisions.

Current understanding of IC afferent organization derives from tracing studies across mammalian species, including cat, ferret, bat, rat, and mouse. Together, these studies established a canonical ascending input pattern to the IC, characterized broadly by predominantly contralateral CN input, as well as bilateral SOC and LL inputs with nucleus-specific differences in laterality (Roth et al., 1978; Beyerl, 1978; Adams, 1979; Brunso-Bechtold et al., 1981; Henkel and Spangler, 1983; Moore, 1988; Bajo et al., 1993; Zhang et al., 1998; Frisina et al., 1998; Kelly et al., 1998; Saldaña et al., 2009; Cant, 2013; Williams and Ryugo, 2024; Rincón et al., 2024). However, in many of these studies, tracer injections spanned large portions of the IC or were centered within the CIC, biasing the canonical pattern toward this core subdivision. Only a few studies compared IC subdivisions (Coleman and Clerici, 1987; González-Hernández et al., 1996; Chen et al., 2018), and even these studies typically used this broad core-versus-shell framework or lacked quantitative analysis. As a result, potential differences between the ECIC and DCIC have remained largely unexplored.

Recent physiological work further suggests that the shell subdivisions should not be treated as a single class when considering ascending inputs. Contrary to the classical view of these regions as slow, modulatory routes, recent studies have reported sound response latencies in the ECIC comparable to those in the CIC (Offutt et al., 2023; Garcia et al., 2025). In both subdivisions, response latencies vary along the rostrocaudal axis, and the rostral ECIC relays short-latency auditory signals to deep layers of the auditory cortex through the non-lemniscal thalamus. These findings make the distinction between the primary CIC and secondary ECIC less clear and suggest that fast ascending pathways also pass through the ECIC in parallel with the canonical CIC pathway. Resolving whether these short-latency responses reflect distinct ascending input patterns within the ECIC therefore requires separate sampling of rostral and caudal subdomains.

In the present study, we performed retrograde tracing from distinct IC subdivisions in mice to systematically characterize their afferent organization. Through selective targeting of the CIC, DCIC, and both rostral and caudal ECIC, and by registering labeled neurons across source nuclei to a publicly accessible brain atlas (Chon et al., 2019; Wang et al., 2020), we directly compared ascending input landscapes across these subdivisions. Our results show that the DCIC diverges substantially from the CIC across major ascending pathways, whereas the ECIC is distinct and does not conform to either pattern. Instead, the ECIC combines CIC-like and DCIC-like inputs in a pathway-dependent manner, forming a hybrid organization rather than a uniform shell identity. In addition, rostral and caudal ECIC exhibit overlapping but distinct input patterns, suggesting further specialization within the ECIC itself. Together, these findings reveal heterogeneity across the shell IC that is not captured by the traditional core-versus-shell framework and provide an anatomical foundation for understanding how parallel midbrain pathways may differentially contribute to auditory processing.

## Materials and Methods

### Animals

Mice were at least 6 weeks old at the time of experiments. The strains used include: C57BL/6J (JAX000664), B6N-Cdh23^tm2.1Kjn^/Kjn (JAX 018399), Sst^tm2.1(cre)Zjh^/J (JAX 013044), Pvalb^tm1(cre)Arbr^/J (JAX 017320), Vip^tm1(cre)Zjh^/J (JAX 010908), Emx1^tm1(cre)Krj^/J (JAX 005628), and Grin1^tm2Stl^/J (JAX 005246). Of the 18 mice (22 injections) included in this study, five animals (six injections) were reanalyzed from our previous study (Garcia et al., 2025). Both female and male mice were used. Animals were housed at 21°C and 40% humidity, with ad libitum access to food pellets and water. They were kept under a reversed light cycle (12–12 h), and all experiments were conducted during their dark cycle. All experimental procedures were approved and conducted in accordance with the Institutional Animal Care and Use Committee at the University of North Carolina at Chapel Hill (protocol 25-124) and the guidelines of the National Institutes of Health.

### Retrograde Tracing

Mice were anesthetized with 4% isoflurane vaporized in 100% oxygen and secured in a stereotaxic frame (Kopf Instruments, Model 1900). Anesthesia was maintained at 1–2% isoflurane throughout the procedure, and body temperature was kept at 35–36.5°C via a feedback-controlled heating pad. Prior to craniotomy, mice received dexamethasone (2 mg/kg body weight) and meloxicam (5 mg/kg) subcutaneously. Enrofloxacin (10 mg/kg) was administered postoperatively before returning mice to a clean cage.

For retrograde tracing from the CIC, DCIC, and caudal ECIC, we visualized the structure by thinning the skull over the IC using a dental drill while intermittently washing the field with phosphate-buffered saline (PBS) to prevent overheating. The IC boundaries were identified as a white rhombus-shaped field lacking surface vasculature and bordered rostro-laterally by the transverse sinus. Once the IC was visualized, the injection sites were determined based on surface landmarks and stereotaxic coordinates. Lateral distances from the midline were approximately 300 µm for DCIC, 1000–1100 µm for CIC, and 1500 µm for caudal ECIC. For caudal ECIC targeting, the head was rolled 24° such that the left ear was elevated above the right while other axes were maintained. Following a small craniotomy, 0.5% fluorophore-conjugated wheat germ agglutinin (WGA-488, 594, or 647) solution was injected through beveled glass pipettes (20 µm external diameter; 20–40 nL at 10–15 nL/min). Injection depths were 600 µm for DCIC, 1000–1200 µm for CIC, and 600–900 µm for caudal ECIC (angled at 24°). Rostral ECIC was stereotaxically targeted at the following coordinates relative to bregma: anterior −4.6 mm, lateral 1.5 mm, ventral 2.35 mm. In two animals, two IC subdivisions were quantified with tracers conjugated to different fluorophores. After injections, the scalp was closed with tissue adhesive, and the mice were recovered from anesthesia before being returned to their home cage.

### Histology and Image Preparation

Three to four days after injection, mice were deeply anesthetized with isoflurane and transcardially perfused with PBS, followed by 4% paraformaldehyde in PBS. Brains were extracted, fixed overnight in 4% paraformaldehyde in PBS, cryoprotected in 30% sucrose in PBS, and coronally sectioned at 40 μm thickness using a freezing microtome. Every third section (approximately 120 µm apart), spanning roughly 3.2 to 7.0 mm posterior to bregma, was mounted on glass slides, counterstained with DAPI, and imaged using an optical sectioning fluorescence microscope (Zeiss Axio Observer 7). In mice used to characterize the dorsal column nuclei, sections were collected up to 9.0 mm posterior to bregma.

WGA has been reported to exhibit transsynaptic anterograde transport under certain conditions. However, transsynaptic labeling typically requires a longer time than what was used here. Consistent with this, at three to four days post-injection, we did not observe anterograde labeling of cell bodies in known downstream target regions, such as the ventral division of the medial geniculate body, following injections in the CIC. This observation supports the interpretation that labeling primarily reflects first-order retrograde transport.

### Atlas Registration and Cell Counting

Whole-brain section images from individual animals were imported into QuPath (v0.5.1; https://qupath.github.io/) and organized as project files (Bankhead et al., 2017). Kim’s Unified Mouse Brain Atlas (https://kimlab.io/brain-map/atlas/) (Chon et al., 2019) and Allen Common Coordinate Framework (CCFv3 2017) (Wang et al., 2020) were obtained through the BrainGlobe Atlas API (https://brainglobe.info/) (Claudi et al., 2020). Coronal sections were registered to the atlas (10 µm resolution) using the Aligning Big Brains and Atlas (ABBA) plugin (v0.11.0; https://abba-documentation.readthedocs.io/en/v0.11.0/) (Chiaruttini et al., 2025). Sections were first positioned along the anterior-posterior (Z) axis at approximately 120 µm intervals. To account for differences in the sectioning angle, the reference atlas was manually rotated along the left-right (X) and dorsal-ventral (Y) axes, using fiber tracts and other anatomical landmarks as guides. The same rotation parameters were applied uniformly across all sections within each project.

We next applied a combination of automated and manual affine and spline transformations to achieve accurate alignment of tissue sections to the reference atlas. Local refinements of anatomical boundaries were performed manually using BigWarp (Bogovic et al., 2016). For the cochlear nucleus, boundaries between the dorsal (DCN) and ventral (VCN) subdivisions were identified primarily based on DAPI staining. Boundaries within the superior olivary complex were determined based on autofluorescence. For the lateral lemniscus, boundaries were determined using a combination of both DAPI staining and autofluorescence to identify the fiber bundle.

Once the transformation from brain tissue sections to the reference atlas was established, transformation parameters were exported back to QuPath for subsequent analysis. We manually annotated presynaptic neurons as detections based on the following criteria: mean fluorescence intensity exceeding the background, an appropriate soma size, clearly defined and non-fragmented morphology, and the presence of visible nuclei stained with DAPI. QuPath recorded the number of detections within each annotated brain region and their spatial coordinates in reference to the Allen Common Coordinate Framework.

### Atlas Curation and Region Definitions

We made several adjustments to the structure tree of Kim’s Unified Mouse Brain Atlas. First, the midbrain raphe nuclei were incorrectly labeled with the acronym “Ramb,” which corresponds to the retroambiguus nucleus. This was corrected to “MBRN,” and the parent acronyms of its child structures were updated accordingly. Second, the corticospinal tract and the commissural stria terminalis shared the same acronym, “cst”. To resolve this ambiguity, the latter was relabeled as “cst2.” Third, the ventral cochlear nucleus granule cell layer (VCAGr) was defined in the structure tree but was not present in the annotated atlas. Instead, a presumably corresponding region was labeled as the granule cell layer of the cochlear nuclei (“GrC (VCCap in 81)”). We renamed this region “VCAGr,” which we consider a more appropriate designation. Fourth, the parent regions of the caudoventrolateral (CVL) and rostroventrolateral (RVL) reticular nuclei, as well as the rostral ventral respiratory group (RVRG), were incorrectly listed as “PGRNl”; we corrected the parent assignment to “MY-mot.” Finally, the name “perilemniscal nucleus, ventral part” (PLV) was inherited from the Paxinos Brain Atlas (George Paxinos, 2012) and likely reflects a misnomer. This was revised to “paralemniscal nucleus, ventral part”. In addition, for consistency with historical and widely used anatomical nomenclature, we refer to the ventral cochlear nucleus anteroventral division (VCA) as the anteroventral cochlear nucleus (AVCN), the ventral cochlear nucleus posteroventral division (VCP) as the posteroventral cochlear nucleus (PVCN), and the dorsal cochlear nucleus (DC) as the DCN throughout the manuscript.

Several regions were combined to simplify analysis because lower hierarchical subdivisions were not consistently distinguishable and some regions contained very low cell counts, making finer parcellation unreliable and difficult to interpret. Lower hierarchical levels were therefore excluded for the following regions: cerebellum (CBL), cerebellar cortex (CBX), dorsal cochlear nucleus (DCN), nucleus raphe (NR), laterodorsal tegmental nucleus (LDT), principal sensory trigeminal nucleus (Pr5), parabrachial nucleus (PB), Barrington nucleus (Bar), dorsal tegmental nucleus (DTg), central gray (CG), reticulotegmental nucleus of the pons (RtTg), motor trigeminal nucleus (5N), raphe magnus nucleus (RMg), solitary nucleus (Sol), abducens nucleus (6N), facial nucleus (7N), ambiguus nucleus (Amb), inferior olivary nucleus (IO), intermediate reticular nucleus (IRt), lateral reticular nucleus (LRt), paramedian reticular nucleus (PMn), parvicellular reticular nucleus (PCRt), lateral paragigantocellular nucleus (LPGi), Botzinger complex (Bo), perihypoglossal nuclei (PHY), vestibulocerebellar nucleus (VeCb), matrix region of the medulla (Mx), superior colliculus (SC), ventral tegmental area (VTA), retroisthmic nucleus (RIs), periaqueductal gray (PAG), interstitial nucleus of Cajal (InC), pretectal region (PRT), cuneiform nucleus (CnF), red nucleus (R), Edinger-Westphal nucleus (EW), trochlear nucleus (4N), substantia nigra, compact part (SNC), pedunculotegmental nucleus (PTg), midbrain raphe nuclei (MBRN), cerebellum-related fiber tracts (cbf), medial forebrain bundle system (mfbs), extrapyramidal fiber systems (eps), and lateral forebrain bundle system (lfbs).

Several regions were further merged to facilitate analysis. First, subdomains of the spinal trigeminal nucleus (Sp5O, Sp5I, and Sp5C) were combined into a single Sp5 region. Second, the posterior ventral cochlear nucleus octopus cell area (VCPO) was merged with the PVCN. Third, the lateral (LVPO) and medial (MVPO) ventral periolivary regions were combined into a single ventral periolivary region (VPO). Fourth, VCAGr was merged with the ventral cochlear nucleus, anterior part (VCA). Finally, the triangular nucleus of the lateral lemniscus (TrLL) was merged with the paralemniscal nucleus (PL).

Inputs from the cerebral cortex (CTX), inferior colliculus (IC), and nucleus of the brachium of the inferior colliculus (BIC) were excluded from quantification to restrict analysis to ascending inputs. For visualization of non-auditory input sources in Fig. 9, reticular regions adjacent to major auditory structures, including the mesencephalic reticular formation (mRt), pontine nuclei (Pn), pontine reticular nucleus (oral part, PnO; caudal part, PnC), and prosomere 1 reticular formation (p1Rt), were excluded to minimize potential misassignment of cells due to ambiguous boundaries.

### Tracer Spread Quantification around Injection Sites

The tracer spread around each injection site was quantified using unsaturated TIFF images. In animals with multiple tracer injections, analyses were performed separately for each fluorescence channel. Images were first smoothed using a Gaussian filter (σ = 10 µm). The maximum fluorescence intensity was identified across all sections. Background for each section was estimated from the image intensity distribution as the average of the median pixel value and the mode of a smoothed histogram. After background subtraction, a tracer spread mask was defined as the contiguous region with intensity greater than 3% of the peak signal across sections. Signal intensity was then quantified from the raw, unsmoothed images, following background subtraction. Boundaries of IC subdivisions were exported from QuPath as ROI polygons in GeoJSON format. Tracer signal was calculated as the summed intensity within the overlap between each ROI and the tracer spread mask. Both hemispheres were included to account for potential contralateral spread. For each injection, signal intensity within each IC subdivision was normalized to the total signal within the tracer spread mask across all sections. Note that this quantification likely overestimates tracer spread due to anterograde and retrograde transport associated with intra-IC connectivity.

### Probability Distributions of Input Neurons

To quantify the spatial distribution of input neurons, we constructed three-dimensional probability distribution maps from cell coordinates exported from QuPath following atlas registration.

Coordinates were converted to voxel indices at 25 µm resolution and mapped onto the annotated brain volume of Kim’s Unified Mouse Brain Atlas. To quantify the spatial probability distribution within individual source structures (LL, SOC, and CN), only input cells located within 0.1 mm of the respective structure boundaries were included. For each animal, voxel-wise three-dimensional cell counts were smoothed with a Gaussian kernel (σ = 75 µm) and normalized to the total cell counts within the corresponding source structure. Coronal slices at selected anteroposterior levels were extracted from the animal-averaged volume and visualized as heat maps overlaid with atlas boundaries.

For visualization of individual neurons, cells within a 200 µm anteroposterior volume were overlaid onto the brain atlas at the center of that volume.

### Similarity Index of Input Patterns

To quantify the similarity of input organization across IC subdivisions, we constructed, for each IC subdivision, an input vector representing the fractional contributions from source structures, with ipsilateral and contralateral inputs concatenated. The similarity index between pairs of IC subdivisions was then calculated as the cosine similarity between their respective input vectors.

For LL inputs, source structures included the dorsal (DLL), intermediate (ILL), and ventral (VLL) nuclei, as well as the paralemniscal nucleus (PL) and its ventral part (PLV), medial paralemniscal nucleus (MPL), and lateral lemniscus fiber tract (*ll*). For SOC inputs, source structures included the lateral (LSO) and medial (MSO) superior olives, dorsal (DPO) and ventral (VPO) periolivary regions, superior paraolivary nucleus (SPO), and nucleus of the central acoustic tract (CAT). For CN inputs, source structures included the DCN, AVCN, and PVCN. For non-auditory inputs, all non-auditory regions contributing at least 0.1% of the total input (71 regions) were included. For total input analysis (Fig. 10E), all brain regions with detected cells in at least three animals (114 regions) were included.

### Statistical analysis

No statistical method was used to predetermine sample size, and sample sizes were determined based on standards commonly used in anatomical tracing studies. All *n* values refer to the number of animals, and key findings were replicated across multiple mice. Mice were randomly assigned to experimental groups. Blinding during surgeries was not feasible, as experimental conditions were defined by anatomical targeting. However, data analysis was performed using predefined criteria. Data are presented as individual data points along with the mean, as specified in the figure legends. Data in texts are presented as mean ± SEM. Statistical comparisons between IC subdivisions were performed using one-way ANOVA followed by Tukey’s honestly significant difference (HSD) post hoc test.

## Results

### Core and shell subdivisions of the IC receive overlapping and distinct ascending inputs

To determine ascending input patterns to individual IC subdivisions, we performed anatomical retrograde tracing experiments. Wheat germ agglutinin conjugated to Alexa Fluor (WGA-488, 594, or 647) was injected into the right CIC, DCIC, and either rostral or caudal ECIC (Fig. 1A–B) (CIC: *n* = 5 mice; rostral ECIC: *n* = 4; caudal ECIC: *n* = 5; DCIC: *n* = 4). The boundary between rostral and caudal ECIC was defined at -4.90 mm from bregma based on previous physiological recordings (Garcia et al., 2025). In some animals, two IC subdivisions were injected with tracers of different fluorophores. After 3–4 days, retrogradely labeled neurons were identified across widely distributed brain regions, including the pons, medulla, midbrain, and cortex (Fig. 1C). To quantify the distribution of input neurons, we aligned the brain section images to the Unified Mouse Brain Atlas (Chon et al., 2019) and registered labeled neurons to individual brain regions. The Unified Atlas integrates the two most widely used mouse brain atlases, the Franklin-Paxinos Brain Atlas (George Paxinos, 2012) and the Common Coordinate Framework (CCF) (Wang et al., 2020), allowing us to combine the CCF-based registration framework with the finer subdivision of auditory structures provided by the Franklin-Paxinos Atlas (Fig. S1A-C).

**Figure 1.**
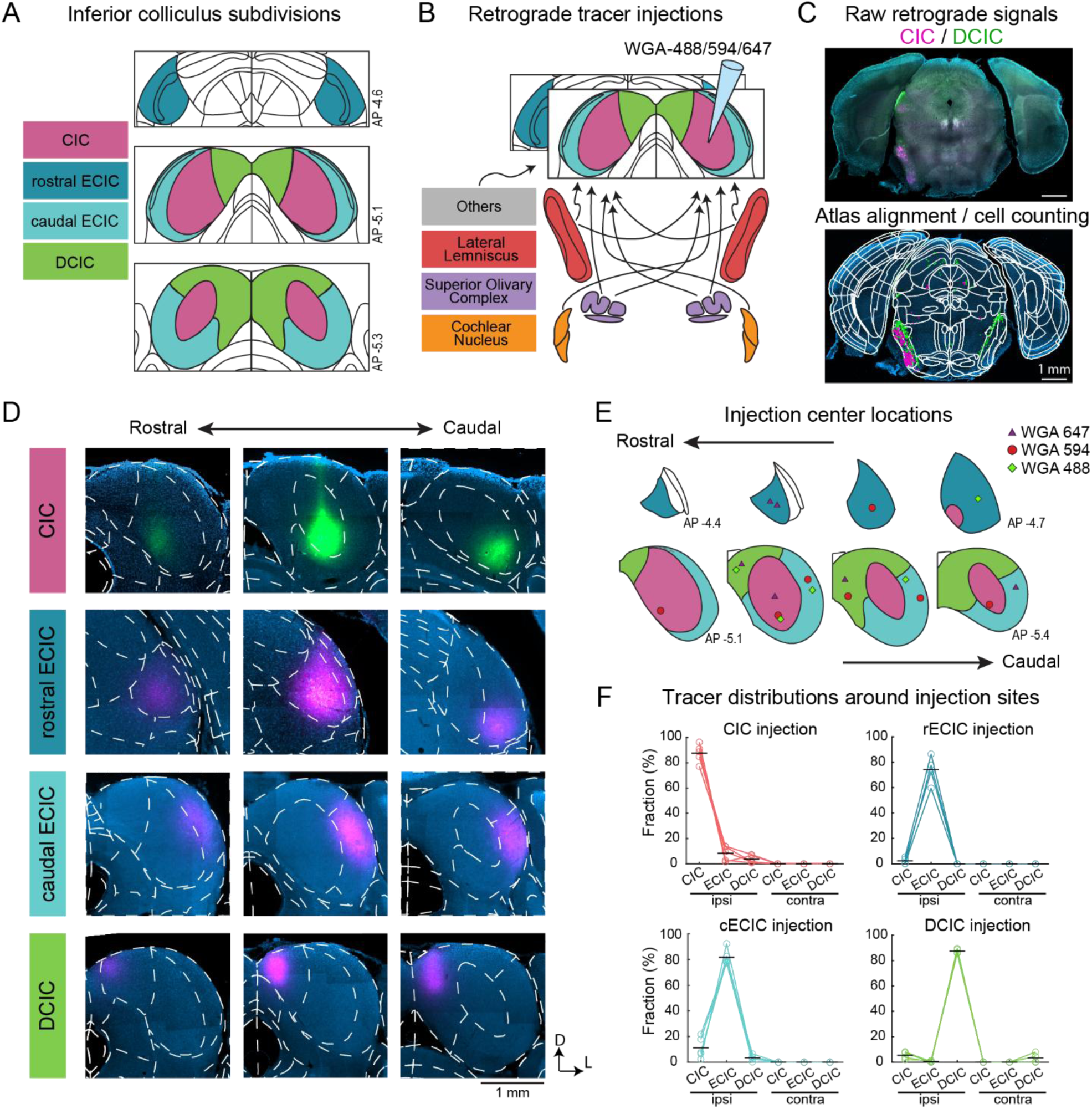
Verification of IC subdivision targeting for retrograde tracer injections. **A,** Schematic diagrams showing IC subdivisions at three anteroposterior planes. **B,** Schematic diagrams illustrating the retrograde tracer injection and major ascending input sources. **C,** Top, raw coronal brain section image showing WGA-594 (red) and WGA-488 (green) signals following injections into the CIC and DCIC, respectively. Bottom, the same section counterstained with DAPI (blue) and overlaid with the registered brain atlas and detected input neurons. **D,** Representative coronal sections of the right IC at three anteroposterior positions around injection sites in the CIC, rostral ECIC, caudal ECIC, and DCIC. **E,** Distribution of injection centers across all mice plotted on the atlas. Different markers represent tracers with distinct fluorophores. **F,** Quantification of tracer spread across IC subdivisions in the ipsilateral and contralateral hemispheres.

We first validated injection targeting by registering the IC to the Unified Atlas and quantifying tracer spread around each injection site (Fig. 1D). Injection center locations within the common atlas space confirmed accurate targeting of the intended IC subdivisions (Fig. 1E). We then quantified the total fluorescence within each brain region and calculated the fraction of tracer signal across subdivisions and hemispheres (Fig. 1F). Because this analysis includes fluorescence from retrogradely labeled somata and anterogradely labeled axons resulting from within-IC connectivity, it likely overestimates cross-subdivisional contamination. Even with this inflated estimate, tracer signals were still largely confined to the targeted subdivisions. For CIC, DCIC, and caudal ECIC injections, mice were included only when >75% of the signal was confined to the target region. In these animals, target confinement averaged 85.5 ± 1.6%, and the greatest cross-subdivisional contamination, observed between the CIC and ECIC, remained below 10% (9.8 ± 2.2%). For DCIC injections, tracer spread to the contralateral DCIC was only 3.3 ± 1.9%, arguing against cross-hemispheric contamination as a confound in analyses of ipsilateral versus contralateral inputs. For rostral ECIC injections, target confinement was slightly lower (74.1 ± 5.6%) due to spread into the brachium of the inferior colliculus (BIC), which overlies the surface of the rostral ECIC. Nevertheless, contamination into other IC subdivisions remained minimal, with the highest spread observed in the CIC at only 2.4 ± 1.3%, indicating negligible cross-subdivisional contamination. Together, these analyses confirm accurate and well-restricted targeting of IC subdivisions for retrograde tracing.

We next examined the spatial distribution of labeled presynaptic neurons across the brain. Representative input cell distributions for CIC, rostral ECIC, caudal ECIC, and DCIC injections reveal clear differences across IC subdivisions even at a macroscopic scale (Fig. 2A–E). For example, strong input from the superior colliculus (SC) was observed in DCIC and ECIC but not in CIC injections. The DCIC showed more bilateral LL input, whereas other IC subdivisions exhibited a stronger ipsilateral bias. In addition, rostral ECIC was unique in showing input from the ventral medullary region extending beyond the CN, including the spinal trigeminal nucleus (Sp5). These observations indicate that IC subdivisions receive distinct patterns of ascending input.

**Figure 2.**
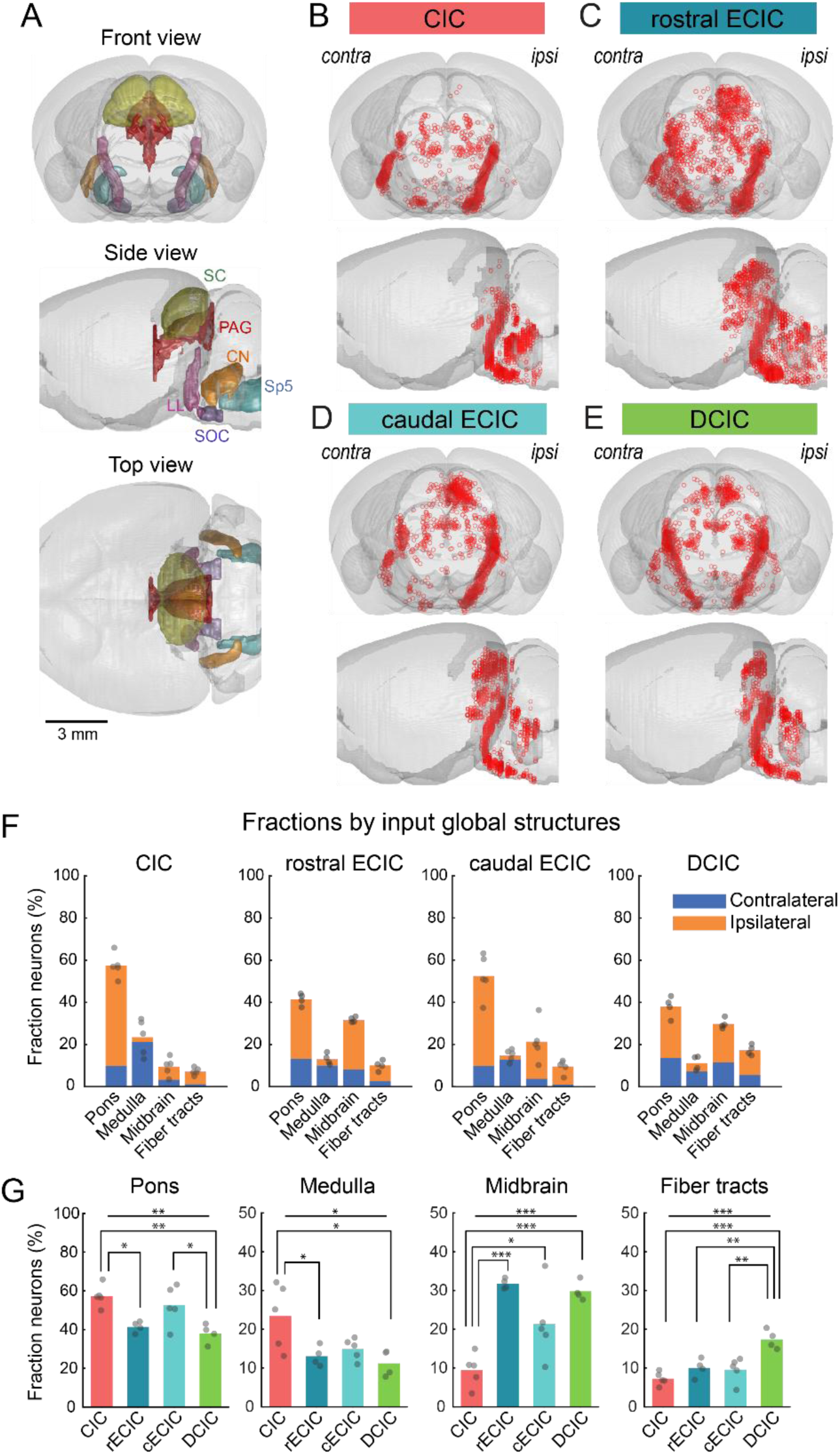
Global-scale distributions of retrogradely labeled neurons across the brain. **A,** Schematic diagrams showing the three-dimensional structures of major source brain regions from the Unified Mouse Brain Atlas, including the whole brain (gray), lateral lemniscus (LL; light purple), superior olivary complex (SOC; dark purple), cochlear nucleus (CN; orange), superior colliculus (SC; green), periaqueductal gray (PAG; red), and spinal trigeminal nucleus (Sp5; light blue). Top, front view; middle, lateral view; bottom, top view. **B–E,** Three-dimensional distributions of presynaptic neurons labeled following retrograde tracer injections into the CIC (B), rostral ECIC (C), caudal ECIC (D), and DCIC (E) in representative mice. Data were subsampled to 2,337 neurons to match cell counts across injection sites (original cell counts: 3,022, 6,566, 2,830, and 2,337, respectively). Top, front view; bottom, side view. **F,** Bar plots showing the fraction of presynaptic neurons in the pons (P), medulla (MY), midbrain (MB), and fiber tracts for injections into the CIC (*n* = 5), rostral ECIC (*n* = 4), caudal ECIC (*n* = 5), and DCIC (*n* = 4). Stacked bars indicate ipsilateral (orange) and contralateral (blue) hemispheres. Individual dots represent individual animals. **G,** Bar plots comparing the fraction of presynaptic neurons across IC subdivisions for each source structure, overlaid with individual animal data. \**p* < 0.05; \*\**p* < 0.01; \*\*\**p* < 0.001 (one-way ANOVA followed by Tukey’s HSD test).

Comprehensive input sources for each injection target, ranked by the total number of labeled neurons across target regions, are provided in Figs. S2–S4.

To further analyze these observations quantitatively, we first grouped ascending source regions into broad anatomical categories: midbrain (MB), pons (P), medulla (MY), and fiber tracts. Across all IC subdivisions, the pons was the largest source of input, reflecting contributions from auditory nuclei, including the LL and SOC, as well as somatosensory/visceral regions such as the principal sensory trigeminal nucleus (Pr5) and parabrachial nucleus (PB), and neuromodulatory regions, including the laterodorsal tegmental nucleus (LDT), locus coeruleus (LC), and raphe nuclei (NR) (Fig. 2F–G; see Fig. 9 for further details).

The second-largest input source distinguished IC subdivisions in a manner broadly consistent with the core-shell dichotomy. In the CIC, the medulla was the second-largest contributor, predominantly reflecting inputs from the dorsal (DCN) and ventral (VCN) cochlear nuclei, with limited input from the midbrain. In contrast, the DCIC and both rostral and caudal ECIC received greater input from the midbrain than from the medulla. Major midbrain inputs to shell subdivisions arose from visual- and motor-related regions, including the superior colliculus (SC), periaqueductal gray (PAG), and cuneiform nucleus (CnF), as well as neuromodulatory regions, including the midbrain raphe nuclei (MBRN) and pedunculotegmental nucleus (PTg). In the DCIC, medullary input was even smaller than that associated with fiber tract regions, including interspersed neurons along the lateral lemniscus (*ll*), cerebellar fiber tracts (cbf), and mesencephalic trigeminal tract (me5). Quantification of fractional input confirmed this distinction; the fraction of medullary input was higher in the CIC than in all other subdivisions (CIC vs. rECIC: *p* = 0.042; cECIC: *p* = 0.085; DCIC: *p* = 0.015; one-way ANOVA followed by Tukey’s HSD test), whereas the fraction of midbrain input was significantly lower in the CIC (CIC vs. rECIC: *p* = 2.3×10^-4^; cECIC: *p* = 0.0238; DCIC: *p* = 5.6×10^-4^) (Fig. 2G). These coarse comparisons provide a basis for the core-versus-shell distinction, which we next resolved at the level of individual input sources.

### Projections from the lateral lemniscus

The lateral lemniscus (LL), located in the pons, constituted the largest ascending input source to the IC. The LL includes the dorsal (DLL), intermediate (ILL), and ventral (VLL) nuclei, together with surrounding paralemniscal regions, including the paralemniscal nucleus (PL), neurons within the lateral lemniscus fiber tract (*ll*), the medial paralemniscal nucleus (MPL), and the ventral part of the paralemniscal nucleus (PLV) (Fig. 3A, C; see Materials and Methods for regional definitions).

**Figure 3.**
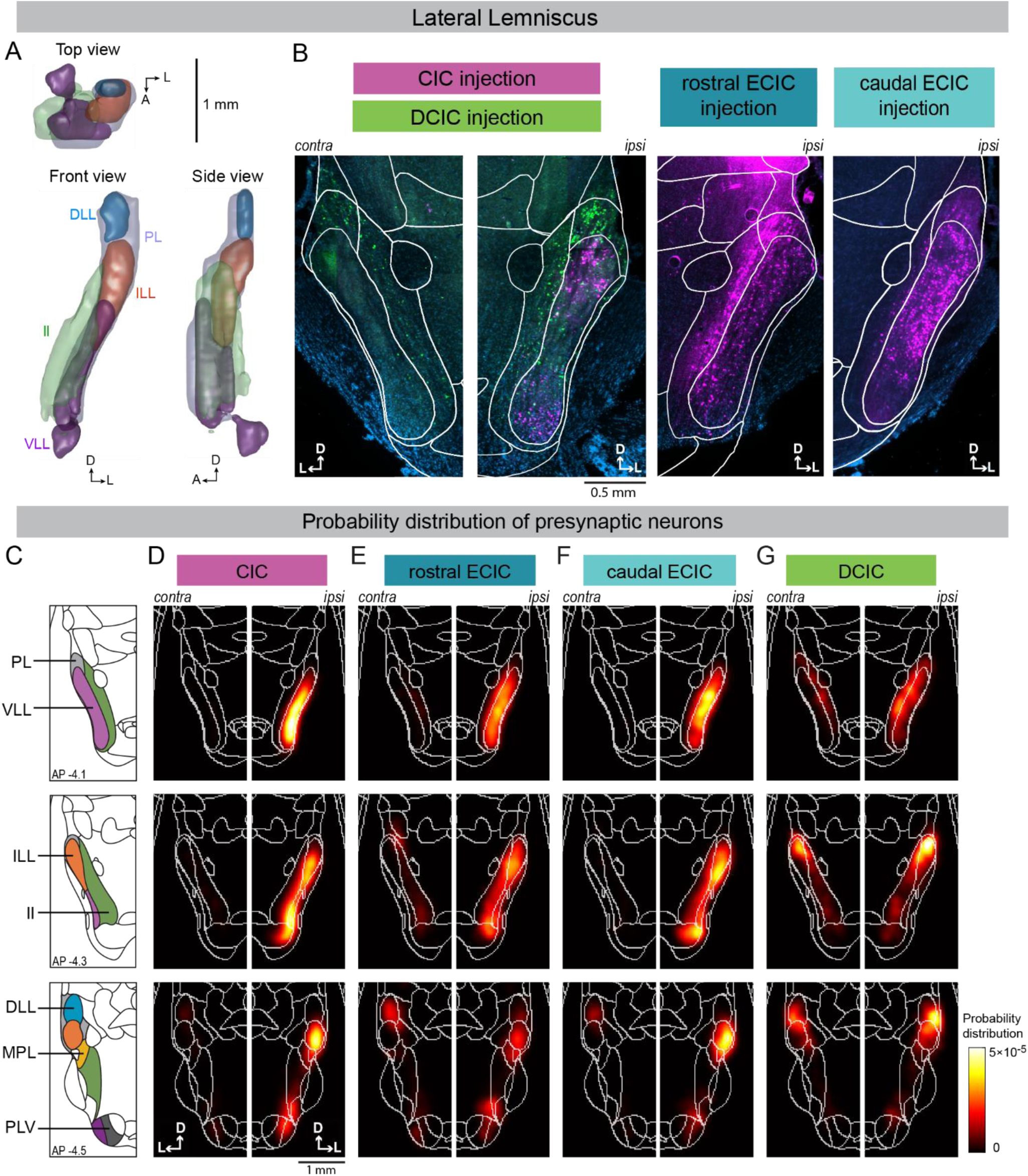
Spatial distribution of projections from the lateral lemniscus. **A,** Three-dimensional structures of regions included in the LL category from the Unified Mouse Brain Atlas, including the DLL (blue), ILL (red), VLL (purple), PL (light blue), and ll (light green), in the ipsilateral hemisphere. Top, front, and side views are shown. **B,** Left, representative coronal sections showing labeling in the LL following injections of WGA-647 (magenta) in the CIC and WGA-488 (green) in the DCIC, displayed for both hemispheres. Middle and right, representative ipsilateral coronal sections from mice injected with WGA-647 in the rostral and caudal ECIC, respectively. Sections were counterstained with DAPI (blue) and overlaid with atlas-derived boundaries. Note that some signals within the ll reflect neuropil labeling rather than somata. **C,** Schematic diagrams showing regional boundaries around the LL at three anteroposterior planes. **D–G,** Probability distributions of retrogradely labeled neurons following injections into the CIC (D), rostral ECIC (E), caudal ECIC (F), and DCIC (G), averaged across animals (*n* = 5, 4, 5, and 4 mice, respectively). Coronal sections correspond to the positions in C.

Retrograde tracer injections into each IC subdivision produced robust labeling in the ipsilateral LL and sparser but reproducible labeling in the contralateral LL. Input patterns differed substantially between IC subdivisions, as illustrated by dual labeling from the CIC and DCIC in the same brain (Fig. 3B). To visualize population-level input patterns, we calculated 3D probability distributions of input neurons within the LL and displayed them as heat maps averaged across animals at representative anterior-posterior positions (Fig. 3C–G). In CIC injections, the strongest labeling was observed in the ipsilateral VLL and ILL, with weaker labeling in the ipsilateral PL and *ll*. Labeling in the DLL was sparse and biased toward the contralateral side (Figs. 3D, 4A). This pattern is consistent with canonical LL projections described in previous bulk-IC tracing studies (Adams, 1979; Brunso-Bechtold et al., 1981; Moore, 1988; Kelly et al., 1998; Zhang et al., 1998). In contrast, DCIC injections produced a distinct labeling pattern, with most labeled neurons located in the ascending fiber tracts of the PL and *ll* (Fig. 3G). Input from the ILL was weak, and labeling in the VLL was much sparser than that observed after CIC injections (Fig. 4A). These paralemniscal-concentrated inputs to the DCIC contrast with those to the CIC, supporting their distinction from the input patterns of the core IC.

**Figure 4.**
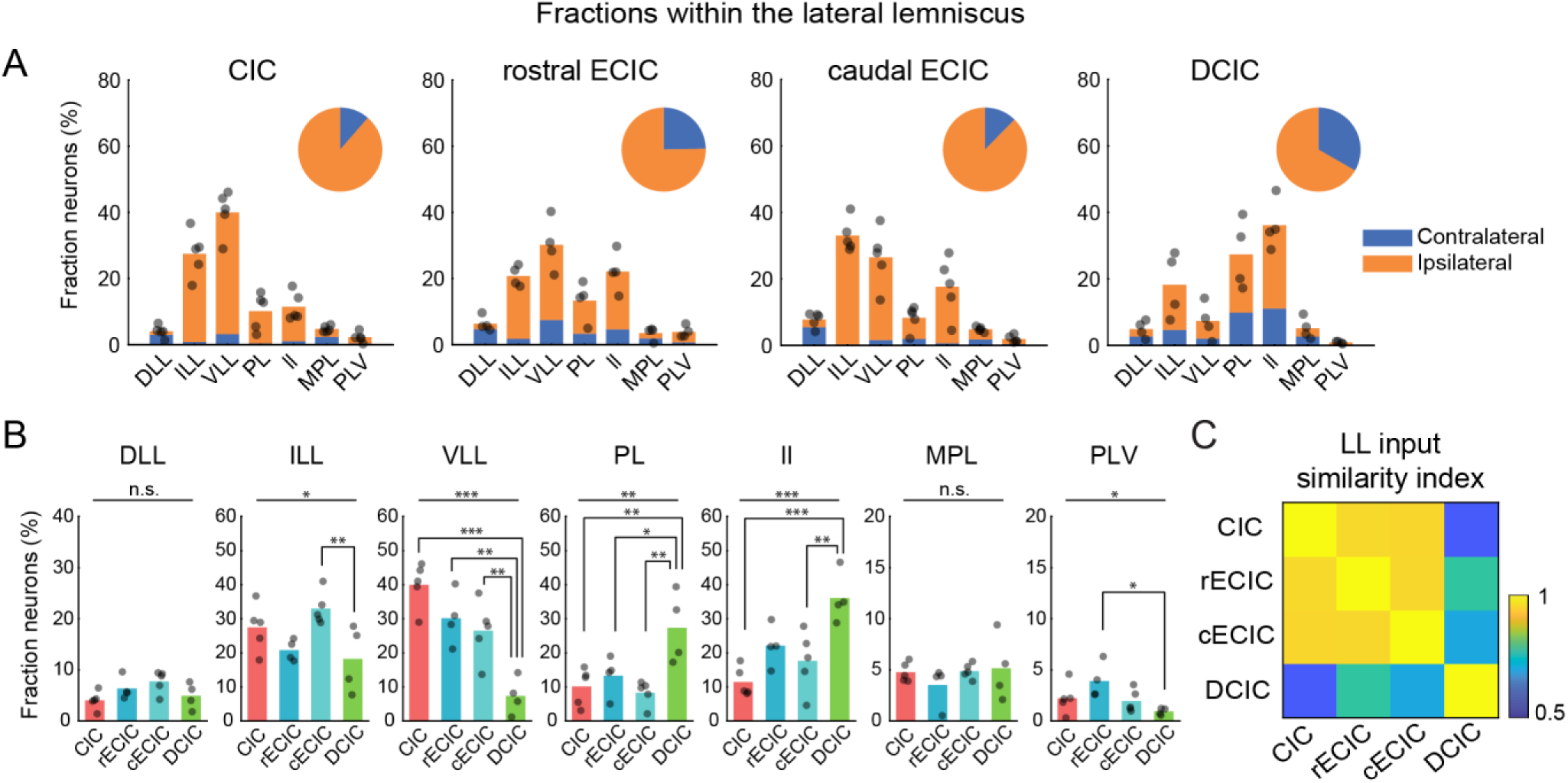
Quantification of projections from the lateral lemniscus. **A,** Bar plots showing the fraction of presynaptic neurons in the DLL, ILL, VLL, PL, ll, MPL, and PLV for injections into the CIC (*n* = 5), rostral ECIC (*n* = 4), caudal ECIC (*n* = 5), and DCIC (*n* = 4). Stacked bars indicate ipsilateral (orange) and contralateral (blue) hemispheres. Individual dots represent individual animals. Inset pie charts show the fractions of ipsilateral and contralateral contributions summed across all LL regions. **B,** Bar plots comparing the fraction of presynaptic neurons across IC subdivisions for each source structure, overlaid with individual animal data. *p < 0.05; **p < 0.01; ***p < 0.001 (one-way ANOVA followed by Tukey’s HSD test). **C,** Heat map showing the similarity index between presynaptic neuron distributions of the CIC, rostral ECIC, caudal ECIC, and DCIC within the LL.

Notably, inputs to both the rostral and caudal ECIC more closely resembled those to the CIC than to the DCIC, although the spatial distributions of labeled cells were somewhat broader than those following CIC injections. In both ECIC subdomains, labeling was strongest in the ILL and VLL, whereas PL- and *ll*-derived inputs were weaker than those after DCIC injections (Figs. 3E–F, 4A). Across animals, the CIC and both ECIC subdomains showed significantly higher fractions of VLL input and significantly lower fractions of PL and *ll* input than the DCIC (Fig. 4B). Both rostral and caudal ECIC also received contralaterally biased input from the DLL, comparable to the CIC. Rostral and caudal ECIC showed broadly similar LL input patterns, with a tendency for a greater ILL contribution in the caudal ECIC.

To quantify these relationships, we calculated the similarity index of fractional input from LL subnuclei across IC subdivisions (see Materials and Methods). Both rostral and caudal ECIC showed greater similarity to the CIC than to the DCIC (Fig. 4C). Together, these results indicate that LL inputs do not follow a simple core-versus-shell dichotomy and instead suggest that the ECIC and CIC share ascending auditory input from the LL.

### Projections from the superior olivary complex

The superior olivary complex (SOC) is another major ascending auditory input source to the IC from the pons and plays a crucial role in sound localization. The SOC includes the lateral and medial superior olives (LSO and MSO), together with surrounding periolivary nuclei, including the superior paraolivary nucleus (SPO), dorsal and ventral periolivary regions (DPO and VPO), and the nucleus of the central acoustic tract (CAT) (Fig. 5A).

**Figure 5.**
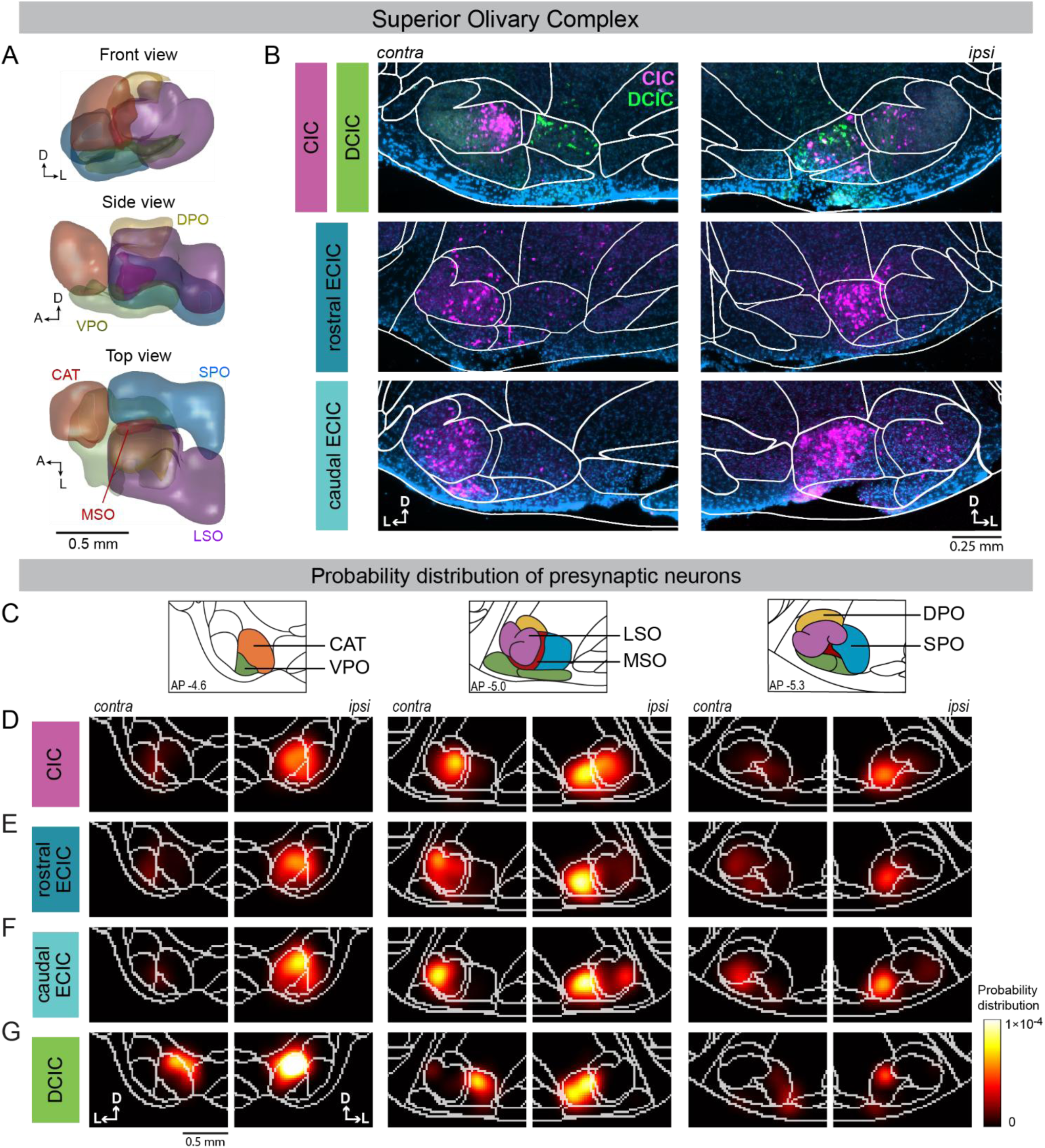
Spatial distribution of projections from the superior olivary complex. **A,** Three-dimensional structures of regions included in the SOC category from the Unified Mouse Brain Atlas, including the LSO (purple), MSO (red), SPO (blue), CAT (orange), DPO (yellow), and VPO (light green), in the ipsilateral hemisphere. Top, front, and side views are shown. **B,** Top, representative coronal sections showing labeling in the SOC following injections of WGA-647 (magenta) in the CIC and WGA-488 (green) in the DCIC, displayed for both hemispheres. Middle and bottom, representative coronal sections from mice injected with WGA-647 in the rostral and caudal ECIC, respectively. Sections were counterstained with DAPI (blue) and overlaid with atlas-derived boundaries. **C,** Schematic diagrams showing regional boundaries around the SOC at three anteroposterior planes. **D–G,** Probability distributions of retrogradely labeled neurons following injections into the CIC (D), rostral ECIC (E), caudal ECIC (F), and DCIC (G), averaged across animals (*n* = 5, 4, 5, and 4 mice, respectively). Coronal sections correspond to the positions in C.

Dual retrograde tracer injections into the CIC and DCIC in the same animal revealed distinct SOC input patterns (Fig. 5B). Population-level probability distributions of input neurons showed that most labeled neurons following CIC injections were located in the contralateral LSO and the ipsilateral CAT and SPO, with a smaller contribution from the VPO (Figs. 5B–D, 6A).

Labeling in the MSO was minimal, consistent with its small size in mice. This overall pattern agrees with previous bulk IC tracing studies, except that CAT input was not characterized in most earlier work (Beyerl, 1978; Roth et al., 1978; Adams, 1979; Brunso-Bechtold et al., 1981; Moore, 1988; Kelly et al., 1998). DCIC injections produced a distinct labeling pattern, with the majority of labeled neurons located bilaterally in the CAT and SPO, though with an ipsilateral bias (Fig. 5G). In contrast, labeling in the LSO and VPO was nearly absent. Thus, inputs to the DCIC were concentrated in the periolivary regions and were distinct from those to the core IC (Figs. 5B, G, 6A).

**Figure 6.**
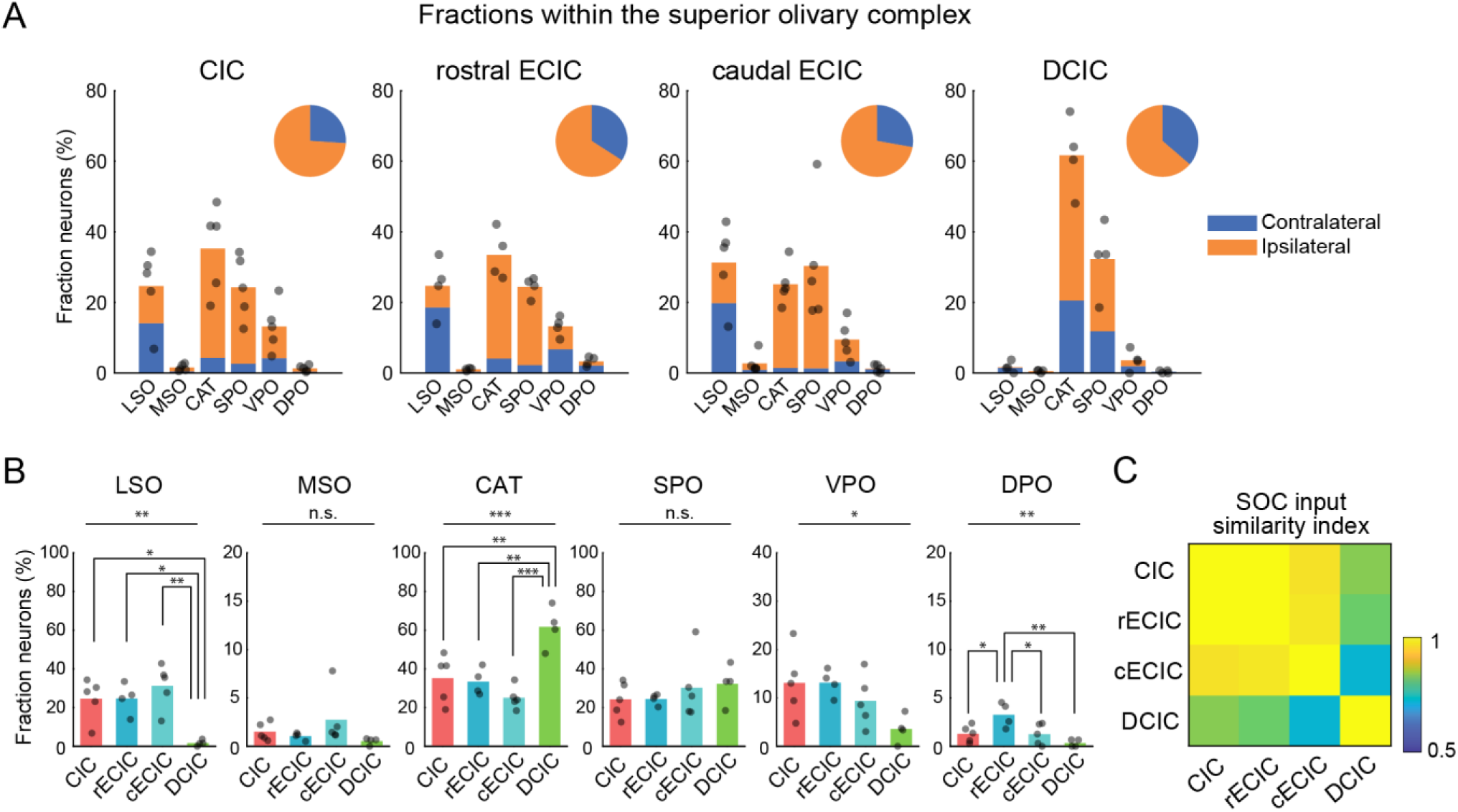
Quantification of projections from the superior olivary complex. **A,** Bar plots showing the fraction of presynaptic neurons in the LSO, MSO, CAT, SPO, VPO, and DPO for injections into the CIC (*n* = 5), rostral ECIC (*n* = 4), caudal ECIC (*n* = 5), and DCIC (*n* = 4). Stacked bars indicate ipsilateral (orange) and contralateral (blue) hemispheres. Individual dots represent individual animals. Inset pie charts show the fractions of ipsilateral and contralateral contributions summed across all SOC regions. **B,** Bar plots comparing the fraction of presynaptic neurons across IC subdivisions for each source structure, overlaid with individual animal data. *p < 0.05; **p < 0.01; ***p < 0.001 (one-way ANOVA followed by Tukey’s HSD test). **C,** Heat map showing the similarity index between presynaptic neuron distributions of the CIC, rostral ECIC, caudal ECIC, and DCIC within the SOC.

SOC inputs to both rostral and caudal ECIC aligned more closely with those to the CIC than to the DCIC. In both ECIC subdomains, labeling remained prominent in the LSO, CAT, and SPO (Figs. 5B,E,F, 6A). Across animals, the CIC and both ECIC subdomains showed significantly higher fractions of LSO input and significantly lower fractions of CAT input than the DCIC (Fig. 6B; Table S1). Thus, input patterns were very similar between the CIC and the two ECIC subdomains except for a small but significantly greater DPO contribution in the rostral ECIC.

To quantify these relationships, we calculated the similarity index using vectors of fractional input from SOC subnuclei across IC subdivisions. Both rostral and caudal ECIC showed greater similarity to the CIC than to the DCIC (Fig. 6C). Together, these results indicate that, similar to the LL inputs, SOC inputs align the ECIC together with the CIC and distinguish both from the DCIC.

### Projections from the cochlear nucleus

Ascending auditory input from the medulla to the IC arises from the cochlear nucleus (CN), the first relay station of auditory information in the brain. The CN comprises the dorsal (DCN) and ventral (VCN) cochlear nuclei, the latter of which can be further divided into anterior and posterior subdivisions (AVCN and PVCN) (Fig. 7A).

**Figure 7.**
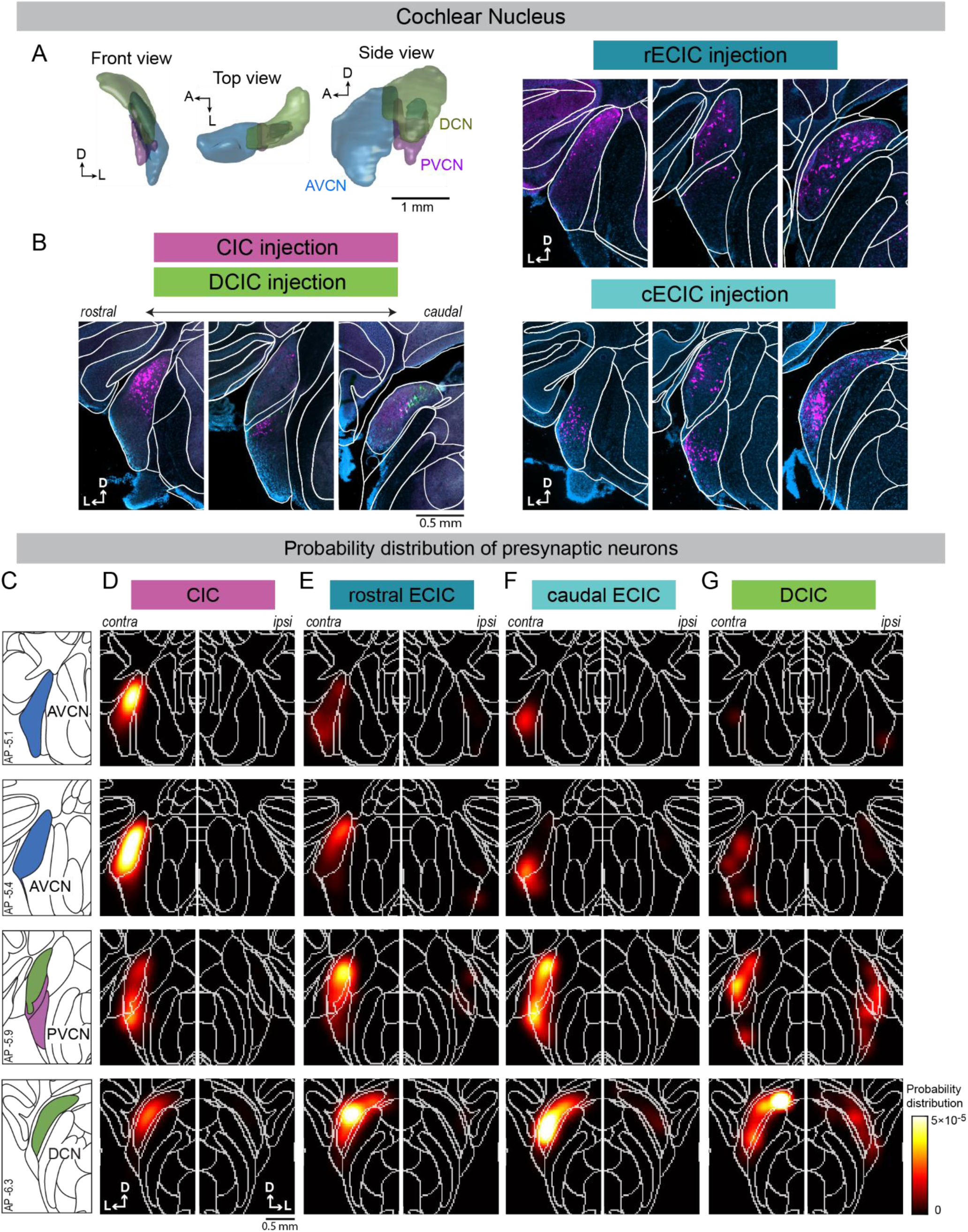
Spatial distribution of projections from the cochlear nucleus. **A,** Three-dimensional structures of regions included in the CN category from the Unified Mouse Brain Atlas, including the AVCN (blue), PVCN (purple), and DCN (green), in the ipsilateral hemisphere. Top, front, and side views are shown. **B,** Left, representative coronal sections showing labeling in the contralateral CN following injections of WGA-647 (magenta) in the CIC and WGA-488 (green) in the DCIC, displayed at three anteroposterior levels. Upper right and lower right, representative coronal sections from mice injected with WGA-647 in the rostral and caudal ECIC, respectively. Sections were counterstained with DAPI (blue) and overlaid with atlas-derived boundaries. **C,** Schematic diagrams showing regional boundaries around the CN at four anteroposterior planes. **D–G,** Probability distributions of retrogradely labeled neurons following injections into the CIC (D), rostral ECIC (E), caudal ECIC (F), and DCIC (G), averaged across animals (*n* = 5, 4, 5, and 4 mice, respectively). Coronal sections correspond to the positions in C.

Dual retrograde tracer injections into the CIC and DCIC in the same animal revealed markedly different CN input patterns (Fig. 7B). Population-level probability distributions showed that CN input to the CIC arose almost exclusively from the contralateral hemisphere (99.5%). Of this input, around 70% originated from the AVCN, with much smaller contributions from the PVCN and DC (Figs. 7B–D, 8A). In contrast, the DCIC received 27% of its CN input from the ipsilateral hemisphere, and most labeled neurons were located in the DCN, with much smaller contributions from the AVCN and PVCN (Figs. 7G, 8A). Thus, CN inputs sharply distinguish the CIC from the DCIC in both laterality and subnuclear origin.

**Figure 8.**
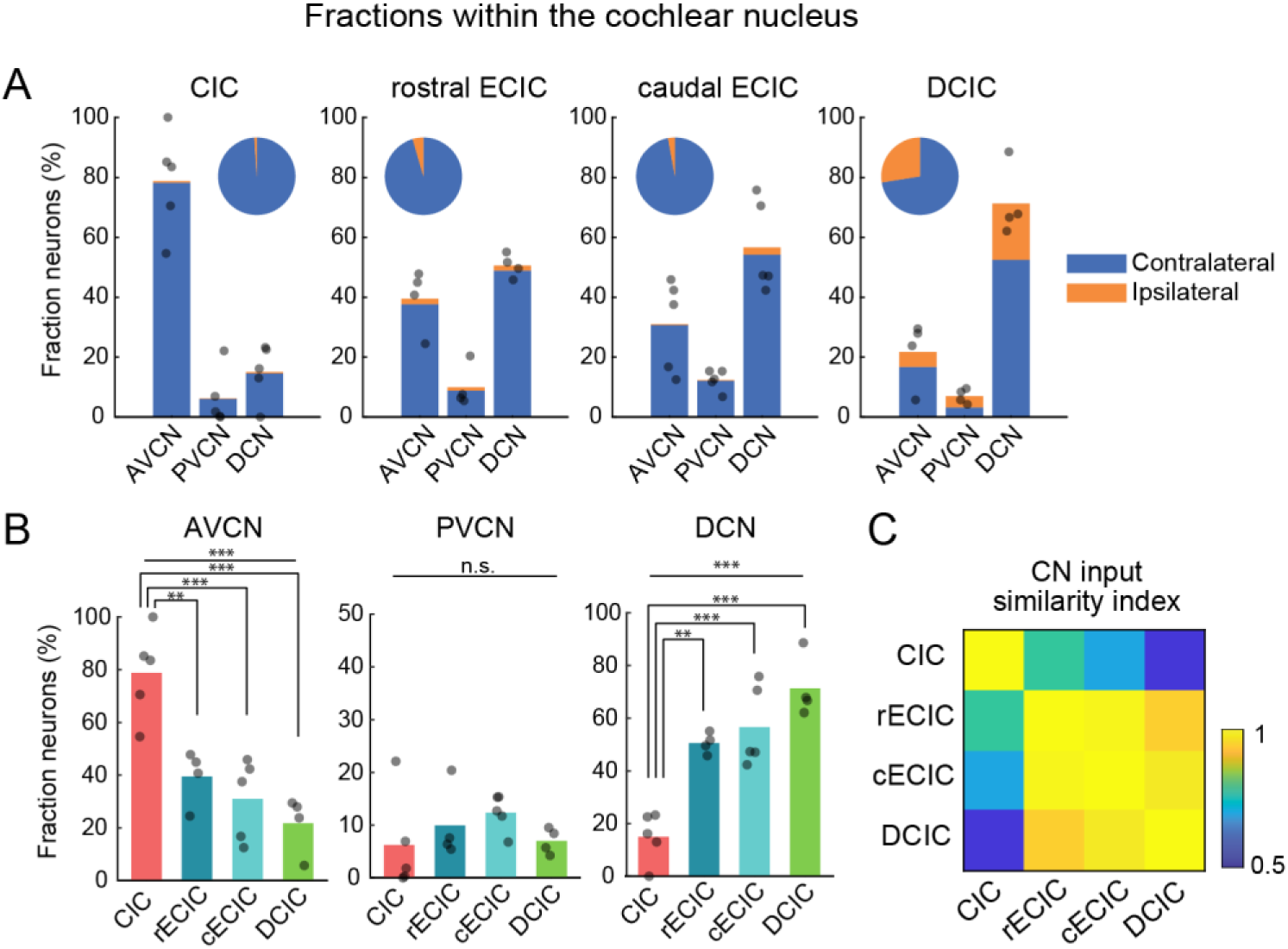
Quantification of projections from the cochlear nucleus. **A,** Bar plots showing the fraction of presynaptic neurons in the AVCN, PVCN, and DCN for injections into the CIC (*n* = 5), rostral ECIC (*n* = 4), caudal ECIC (*n* = 5), and DCIC (*n* = 4). Stacked bars indicate ipsilateral (orange) and contralateral (blue) hemispheres. Individual dots represent individual animals. Inset pie charts show the fractions of ipsilateral and contralateral contributions summed across all CN regions. **B,** Bar plots comparing the fraction of presynaptic neurons across IC subdivisions for each source structure, overlaid with individual animal data. \*\*\**p* < 0.001 (one-way ANOVA followed by Tukey’s HSD test). **C,** Heat map showing the similarity index between presynaptic neuron distributions of the CIC, rostral ECIC, caudal ECIC, and DCIC within the CN.

In contrast to LL and SOC inputs, CN input to the ECIC showed an intermediate pattern but aligned more closely with that to the DCIC (Figs. 7E–F, 8A). Caudal ECIC closely resembled the DCIC, with ∼60% of input originating from the DCN. Rostral ECIC showed a tendency for a larger contribution from the AVCN (rECIC: 40.2%, cECIC: 27.9%), resulting in nearly balanced input from VCN and DCN (VCN: 49.9%; DCN: 50.1%) (Fig. 8A–B). The spatial distribution of labeled neurons within the AVCN following ECIC injections was concentrated near the lateral and medial edges, compared to CIC injections, and neurons labeled from rostral ECIC clustered in the rostrodorsal AVCN (Fig. 7A–B, E). In rostral and caudal ECIC injections, 4.5% and 3.5% of CN input arose from the ipsilateral hemisphere, respectively. These results are consistent with our previous tracing study using cholera toxin subunit B (Garcia et al., 2025).

We next quantified the similarity of CN input patterns across IC subdivisions based on the fractional contributions of CN subnuclei. Both ECIC subdivisions were more similar to the DCIC than to the CIC; however, rostral ECIC showed greater similarity to the CIC than caudal ECIC did (Fig. 8C). Thus, unlike the LL and SOC, CN input places the ECIC closer to the DCIC while also revealing rostro-caudal differences within the ECIC itself.

### Projections from non-auditory brain regions

Finally, retrograde tracing also revealed inputs to the IC from a diverse set of non-auditory regions (Fig. 9). These inputs were not uniformly distributed across IC subdivisions but instead showed an overall bias toward the ECIC and DCIC relative to the CIC.

**Figure 9.**
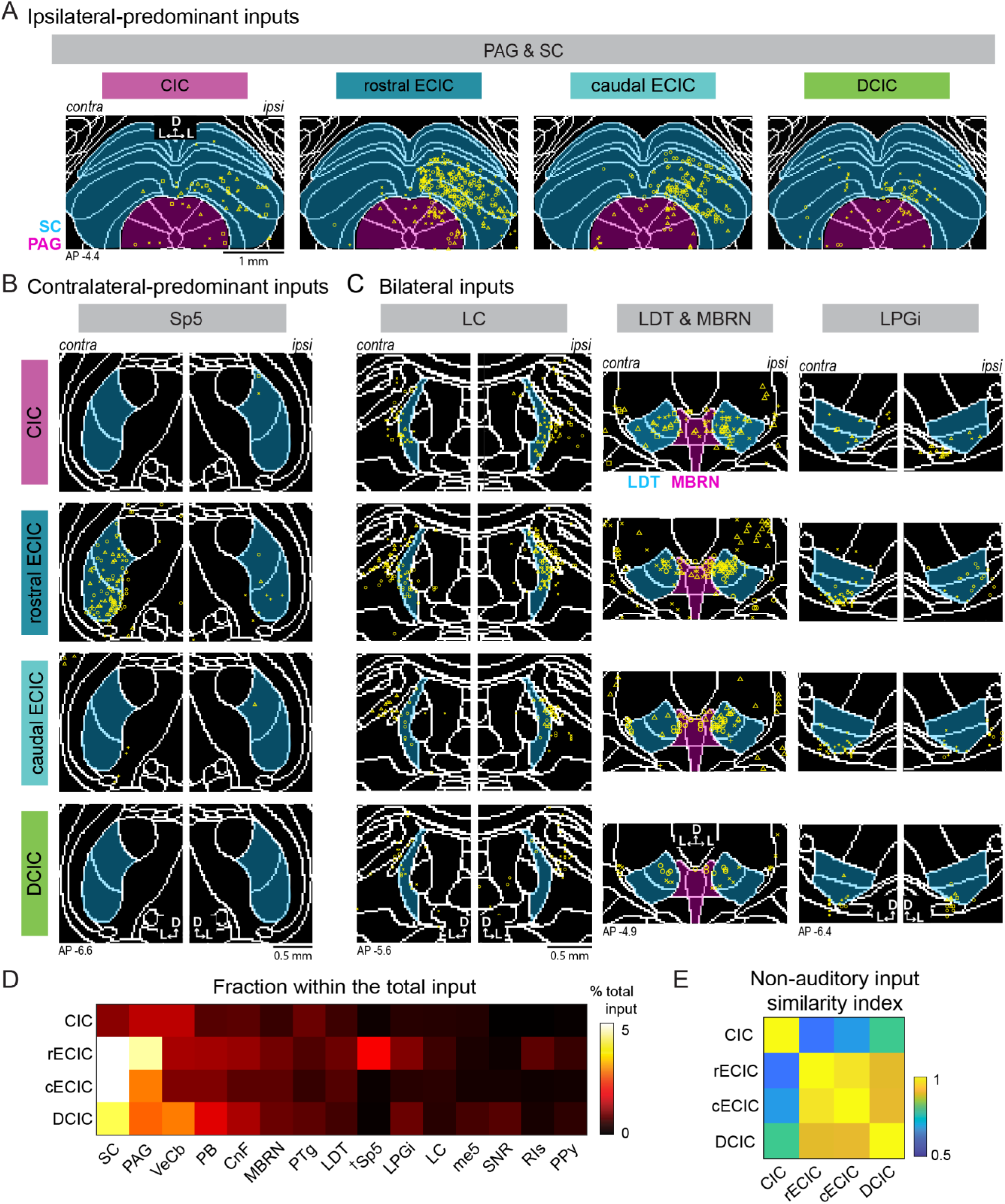
Spatial distribution of projections from non-auditory structures. **A,** Distributions of retrogradely labeled neurons following injections into the CIC, rostral ECIC, caudal ECIC, and DCIC, overlaid across animals (*n* = 5, 4, 5, and 4 mice, respectively), in regions surrounding the SC (blue) and the PAG (magenta). Neurons from different animals are represented by distinct symbols. **B,** Same as in A, for neurons surrounding the Sp5 (blue). **C,** Left, same as in A, for neurons surrounding the LC (blue). Middle, for neurons surrounding the LDT (blue) and MBRN (magenta). Right, for neurons surrounding the LPGi (blue). **D,** Heat map showing the fraction of total input from individual non-auditory sources, averaged across animals. ^†^Quantification for the Sp5 does not cover its full caudal extent. **E,** Heat map showing the similarity index between presynaptic neuron distributions of the CIC, rostral ECIC, caudal ECIC, and DCIC for non-auditory sources.

These sources fell broadly into two categories: crossmodal sensorimotor/visceral regions, including the superior colliculus (SC), periaqueductal gray (PAG), vestibular cerebellar nuclei (VeCb), parabrachial nucleus (PB), cuneiform nucleus (CnF), and spinal trigeminal nucleus (Sp5), and brain state-related regions, including the midbrain raphe nuclei (MBRN), pedunculotegmental nucleus (PTg), laterodorsal tegmental nucleus (LDT), lateral paragigantocellular nucleus (LPGi), and locus coeruleus (LC). We did not include the dorsal column nuclei, a known source of somatosensory input to the ECIC (Björkeland and Boivie, 1984; Coleman and Clerici, 1987; Lesicko et al., 2016; Huey et al., 2025), because these caudal nuclei were poorly covered by the Unified Atlas. Nevertheless, we confirmed their projections to both rostral and caudal ECIC in an additional non-quantitative observation (Fig. S5).

Crossmodal sensorimotor and visceral regions generally showed stronger target selectivity and lateralization than brain state-related sources. The strongest non-auditory inputs arose from the SC and PAG, which both showed preferential projections to the ECIC and DCIC with an ipsilateral bias (SC, IC subdivisions: *p* = 0.0025; CIC vs. rECIC: *p* = 0.0017; CIC vs. cECIC: *p* = 0.0307; PAG, IC subdivisions: *p* = 0.0013; CIC vs. rECIC: *p* = 6.5×10^-4^; rECIC vs. cECIC: *p* = 0.0496; rECIC vs. DCIC: *p* = 0.0368; one-way ANOVA followed by Tukey’s HSD test) (Fig. 9A, D; see Table S1 for full statistics). In contrast, the rostral portion of Sp5 covered by the atlas, comprising the oral (Sp5O) and intermediate (Sp5I) parts, showed highly selective connectivity with the rostral ECIC with a contralateral bias. Labeled neurons were not detected following injections in other subdivisions, including the caudal ECIC (IC subdivisions: *p* = 7.3×10^-7^; rECIC vs. CIC: *p* = 2.6×10*^-6^*; cECIC: *p* = 2.3×10*^-6^*; DCIC: *p* = 3.7×10*^-6^*) (Fig. 9B, D). Qualitative observation of the caudal part of Sp5 (Sp5C), which lies outside the atlas extent, revealed the opposite pattern with selective projections to the caudal ECIC (Fig. S5; see Discussion). Together, these results support preferential targeting of shell subdivisions by crossmodal sources while also revealing input specialization along the rostrocaudal axis of the ECIC.

Brain state-related regions generally provided broader, bilateral, and relatively weaker projections to IC subdivisions. The MBRN, including the dorsal raphe, as well as PTg, LDT, and the LC, projected bilaterally to all IC subdivisions without significant target preference (Fig. 9C–D). One exception was the LPGi, which showed significantly stronger input to the rostral ECIC than to the other subdivisions (IC subdivision: *p* = 0.010; rECIC vs. CIC: *p* = 0.0261; cECIC: *p* = 0.0273) (Fig. 9C–D).

To summarize non-auditory input patterns across IC subdivisions, we quantified the similarity index using 71 non-auditory regions that contributed at least 0.1% of total input (Fig. 9E). Both rostral and caudal ECIC showed greater similarity to the DCIC than to the CIC, consistent with a broad core-versus-shell distinction. At the same time, the two ECIC subdomains clustered together and remained distinct from the DCIC, indicating additional specialization within the shell.

Together with the auditory structure analyses above, these data show that IC subdivisions are distinguished not only by canonical auditory inputs but also by distinct patterns of non-auditory connections.

## Discussion

### The ECIC as an interface between CIC-like and DCIC-like ascending inputs

In this study, we systematically mapped ascending inputs to subdivisions of the mouse IC using retrograde tracing combined with atlas-based registration. In contrast to the prevailing core-versus-shell framework, our results reveal substantial heterogeneity in ascending input organization across shell subdivisions of the IC (Fig. 10A–D). The CIC showed an input pattern that closely matched the canonical organization described in classical tracing studies, supporting both the validity of our dataset and the interpretation that the established literature has predominantly described a CIC-dominated ascending pathway (Beyerl, 1978; Roth et al., 1978; Adams, 1979; Brunso-Bechtold et al., 1981; Kelly et al., 1998). The DCIC clearly contrasted with the CIC in its ascending input patterns at all stages of auditory processing, with inputs originating predominantly from paralemniscal and periolivary nuclei. In contrast, the ECIC did not conform to a simple CIC-like or DCIC-like pattern and instead showed a pathway-specific hybrid organization. Inputs from the LL and SOC grouped the ECIC with the CIC, whereas inputs from the CN and non-auditory brainstem structures grouped it more closely with the DCIC (Fig. 10E–F).

**Figure 10.**
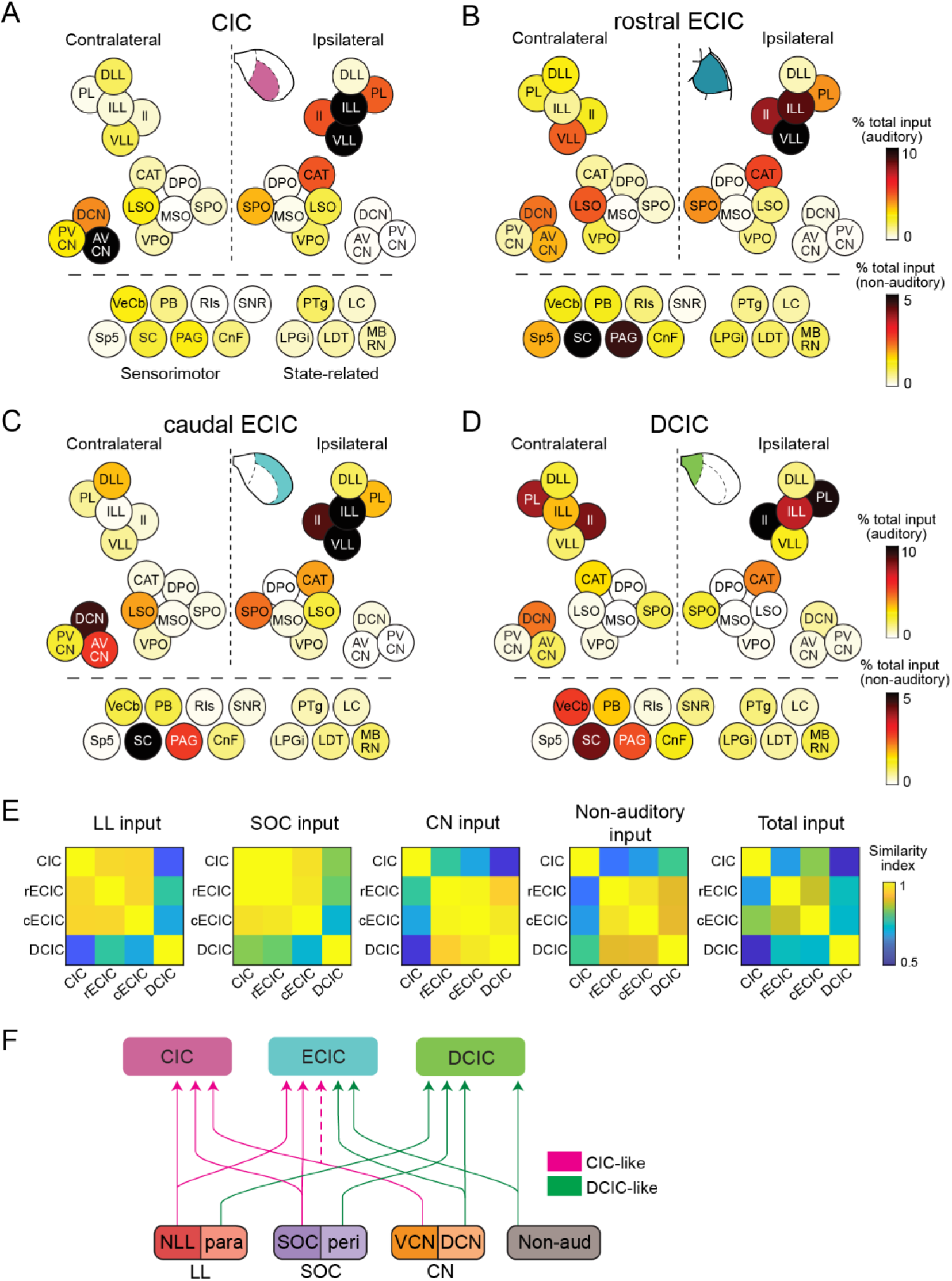
Summary of ascending inputs. **A–D,** Heat maps showing the fraction of total input from individual auditory (top) and non-auditory (bottom) sources for the CIC (A), rostral ECIC (B), caudal ECIC (C), and DCIC (D), averaged across animals and grouped by category. **E,** Heat maps showing the similarity index between presynaptic neuron distributions of the CIC, rostral ECIC, caudal ECIC, and DCIC for the LL (from Fig. 4C), SOC (from Fig. 6C), CN (from Fig. 8C), non-auditory sources (from Fig. 9E), and total input. **F,** Summary schematic illustrating inputs to the CIC, ECIC, and DCIC from auditory and non-auditory sources. CIC-like inputs, predominantly from the nuclei of the lateral lemniscus (NLL), SOC, and VCN, are shown in magenta. DCIC-like inputs, predominantly from paralemniscal regions (para), periolivary regions (peri), DCN, and non-auditory sources, are shown in green. The ECIC exhibits a pathway-specific hybrid organization combining CIC-like and DCIC-like inputs.

Additionally, the ECIC received non-auditory inputs distinct from those of the DCIC and varied along its rostrocaudal axis. Thus, the ECIC is not simply an intermediate between the CIC and DCIC on a single continuum, but rather serves as a hybrid interface at which distinct ascending input motifs converge earlier in the auditory pathway than is typically assumed.

This framework may help explain why ECIC neurons have been associated with both short-latency auditory responses and broader integrative properties (Bajo et al., 2010; Wong and Borst, 2019; Offutt et al., 2023; Quass et al., 2024; Garcia et al., 2025; Ibrahim et al., 2025). CIC-like LL and SOC inputs, together with direct CN input, could provide rapid sound information that can be relayed to higher-order brain regions (Garcia et al., 2025). At the same time, DCIC-like CN and non-auditory inputs, as well as corticofugal input (Winer et al., 1998), would allow the ECIC to participate in the broader multimodal and contextual interactions long associated with non-central IC regions. The ECIC may therefore occupy a functionally advantageous position, in which state, context, or crossmodal signals enable flexible sound processing while maintaining short-latency auditory drives comparable to those of the CIC. Although atlas-based retrograde tracing does not resolve synaptic strength, transmitter identity, and whether distinct source systems converge onto the same IC neurons, it lays the groundwork for testing how auditory, somatosensory, and contextual signals are integrated across ECIC and DCIC microcircuits.

### Potential substrates for fast responses and rostrocaudal specialization in the ECIC

Recent physiological studies have demonstrated that the ECIC exhibits sound response latencies as short as those in the CIC and can relay short-latency auditory signals to higher-order cortical areas (Offutt et al., 2023; Garcia et al., 2025). Our tracing data suggest at least two non-mutually exclusive anatomical substrates for these short-latency responses. First, both rostral and caudal ECIC retained strong LL and SOC inputs shared with the CIC. These primary ascending pathways could therefore convey sound information to the CIC and ECIC at comparable latencies. Second, the rostral ECIC received a greater contribution from the AVCN than the caudal ECIC, suggesting that it may receive short-latency input directly from the AVCN, a known fast auditory relay to the CIC. The concentration of retrogradely labeled neurons in the dorsal AVCN, which represents high frequencies, is also consistent with previous work reporting a high-frequency bias in sound responses in the rostral ECIC (Offutt et al., 2023). Together, these inputs provide an anatomical explanation for fast ECIC responses and suggest that LL/SOC and AVCN inputs constitute two distinct routes for short-latency drive to the ECIC.

Rostrocaudal specialization within the ECIC also extends beyond auditory pathways. We observed a spatial segregation of inputs from the Sp5, a structure associated with somatosensory signals from the facial region (Coleman and Clerici, 1987; Zhou and Shore, 2006). Within the rostral parts of Sp5 covered by the Unified Mouse Brain Atlas, the oral (Sp5O) and intermediate (Sp5I) subdivisions both projected selectively to the rostral ECIC. In contrast, the caudal part (Sp5C), which lies outside the extent of the atlas, projected exclusively to the caudal ECIC (Fig. S5). These Sp5 subdivisions are themselves functionally differentiated: Sp5O is associated with oral and perioral processing, Sp5I shows mixed response properties and broader receptive fields, and Sp5C is more often linked to nociceptive and thermosensory signals from the broader face and upper neck regions. The rostrocaudal segregation of Sp5 inputs therefore suggests that the rostral and caudal ECIC may support distinct forms of somatosensory integration, potentially reflecting differences in somatotopy or sensory modality.

The ECIC is also known to receive somatosensory input from the dorsal column nuclei (the cuneate and gracile nuclei), located at the junction between the medulla and spinal cord (Björkeland and Boivie, 1984; Coleman and Clerici, 1987; Lesicko et al., 2016; Huey et al., 2025). We did not include the dorsal column nuclei in the quantitative analysis because this caudal medullary region is poorly represented in the Unified Atlas. Nevertheless, in a separate qualitative assessment, we observed labeled neurons in the dorsal column nuclei after both rostral and caudal ECIC injections, but not after CIC or DCIC injections (Fig. S5), consistent with previous reports. Together, these observations suggest that both rostral and caudal ECIC may receive overlapping vibratory and other body-related somatosensory input via the dorsal column nuclei, while differing in their trigeminal/facial somatosensory inputs.

The organization of somatosensory input-recipient GABAergic “modules” is also consistent with rostrocaudal distinction: in the caudal ECIC, these modules appear as isolated islands, whereas rostrally they merge to form a more continuous lamina (Lesicko et al., 2016, 2020). However, we note that our tracer injections were not designed to distinguish module from matrix compartments or superficial from deep layers. The ECIC also exhibits tonotopy along its mediolateral (dorsoventral) axis, but we restricted CIC and ECIC injections to their mid-to-deep portions to minimize spread into neighboring subdivisions. The present data therefore cannot determine whether the pathway-specific hybrid pattern we describe is distributed broadly across the ECIC or instead attributable to distinct microdomains within it. Determining how the differential connectivity identified here maps onto modules, matrices, layers, and tonotopic domains within the ECIC will be an important direction for future work.

### Comparison with previous tracing studies

The ascending input pattern to the CIC in our study was largely consistent with canonical ascending pathways described in earlier tracing studies across cat, rat, ferret, and mouse, including strong contralateral CN input, bilateral LSO and DLL input, and ipsilateral input from the ILL, VLL, MSO, and SPO (Roth et al., 1978; Beyerl, 1978; Adams, 1979; Brunso-Bechtold et al., 1981; Henkel and Spangler, 1983; Moore, 1988; Bajo et al., 1993; Zhang et al., 1998; Frisina et al., 1998; Kelly et al., 1998; Saldaña et al., 2009; Cant, 2013; Williams and Ryugo, 2024; Rincón et al., 2024). Fewer studies have examined ascending inputs to the shell subdivisions in comparable detail (Coleman and Clerici, 1987; González-Hernández et al., 1996; Chen et al., 2018). Although prior work generally agreed that the IC core and shell differ in their afferent inputs, these studies typically combined the ECIC and DCIC or did not provide quantitative analysis. Our results therefore extend the earlier literature by providing, to our knowledge, the first systematic quantification of subdivision-specific ascending input patterns to the ECIC and DCIC and by showing that these two shell subdivisions are themselves clearly distinct.

The present study highlights several input sources that were not separately resolved or emphasized in many earlier retrograde studies, including the CAT, PL, and neurons associated with the *ll* fiber tract. One particularly notable feature of our dataset is the prominent input from the CAT to all IC subdivisions. This region, located at the transition between the SOC and LL, was not included in the quantification of most earlier studies, with a rare exception in bats (Casseday et al., 1989; Behrend and Schuller, 2000). Likewise, the PL and neurons associated with the *ll* fiber tract were among the major contributors to DCIC input. These paralemniscal and periolivary structures were likely underrepresented in earlier work because many studies focused on the major classical nuclei rather than on smaller surrounding nuclei and transition zones. Their prominence here suggests that subdivision-specific computation in the IC depends not only on inputs from the major auditory nuclei, but also on adjacent paralemniscal and periolivary regions, whose functional roles remain poorly understood.

Our results also help reinterpret earlier studies that might otherwise appear discrepant. A previous monosynaptic rabies tracing study concluded that shell IC neurons receive predominantly ascending input, helping shift the field away from the view that shell regions are primarily dominated by descending input (Chen et al., 2018). At the same time, they reported relatively limited differences between core and shell across many of the major ascending auditory source nuclei, aside from stronger VCN input to the core. Although tracing using the rabies virus and WGA may not be directly comparable, our findings suggest a plausible explanation for that result. If the ECIC and DCIC are pooled, the strong LL and SOC inputs shared by the CIC and ECIC would increase the apparent similarity between core and shell. In addition, because paralemniscal and periolivary nuclei were not separately quantified, much of the input that distinguishes the DCIC from both the CIC and ECIC may not have been captured. Our results therefore extend, rather than contradict, Chen et al. by identifying heterogeneity between the ECIC and DCIC.

A few source-specific anterograde tracing studies may appear, at first glance, to conflict with the robust SOC input to the ECIC observed here. In cats and Mongolian gerbils, LSO and MSO projections have been described as terminating within the CIC, including the so-called pars lateralis (Shneiderman and Henkel, 1987; Oliver et al., 1995; Cant, 2013). However, under the cross-species parcellation proposed by Loftus et al. (2008), territory historically labeled pars lateralis in cats corresponds more closely to the deep lateral cortex, making it more comparable to rodent ECIC than to the CIC (Loftus et al., 2008). Under this revised framework, the earlier anterograde tracing data from the LSO and MSO are compatible with the idea that SOC projections are shared by the CIC and ECIC rather than restricted to the CIC alone. Apparent discrepancies with the older literature may therefore reflect differences in subdivision definitions more than genuine disagreement about projection patterns.

Finally, we note two aspects of the Unified Mouse Brain Atlas that may benefit from refinement. First, the region where the auditory nerve enters the CN is labeled as the vestibulocochlear nerve (8n) but is typically considered part of the AVCN and PVCN. We observed many input neurons in this region, supporting its classification as part of the VCN. Second, the ventral part of the VLL appears discontinuous in the 3D-reconstructed atlas volume (Fig. 3A), due to an incursion of the pontine nucleus (Pn). However, our 3D probability distribution maps consistently showed strong IC inputs from this region (Fig. 3D–G), suggesting that it more appropriately belongs to the LL and may represent a continuous extension of the VLL. We therefore did not treat the Pn as a major non-auditory input source, despite its relatively high contribution in the quantified input fractions (Fig. S2). Despite these limitations, the Unified Atlas provides more detailed delineation of brainstem auditory nuclei than the Allen CCF (Fig. S1) and remains a valuable resource for future studies.

Overall, our results refine the classical core-versus-shell framework of the IC by showing that shell subdivisions are not homogeneous: the ECIC combines CIC-like and DCIC-like ascending input motifs, whereas the DCIC exhibits a distinct input organization. Because the rostral ECIC can provide short-latency auditory drive to deep layers of both primary and secondary auditory cortices (Garcia et al., 2025), this pathway may deliver auditory information that has already been shaped by multisensory and contextual signals at the midbrain level. Modulation of this parallel ascending route could therefore support more flexible cortical processing than the canonical CIC-dependent pathway alone.

## Data Availability

Raw brain section images supporting the findings of this study will be deposited in the DANDI Archive and made publicly available upon publication.

## Author contributions

H.C.A., M.R.K., P.B.M., and H.K.K. designed research. H.C.A., M.M.G., and H.K.K. performed research. H.C.A., M.M.G., M.A-M., and H.K.K. analyzed data. H.C.A. and H.K.K. wrote the manuscript with input from all authors.

## Acknowledgments

This work was supported by the National Institute on Deafness and Other Communication Disorders (R01DC017516: H.K.K.) and the National Institute of Neurological Disorders and Stroke, BRAIN Initiative (R01NS128873: H.K.K., P.B.M., and M.R.K.). We thank Hiroaki Tsukano, Gates Schneider, and Kendall Hutson for their advice throughout the project and comments on the manuscript.

