## Supplementary Figures for "Ascending inputs to inferior colliculus subdivisions reveal pathway-specific hybrid organization in the external cortex"

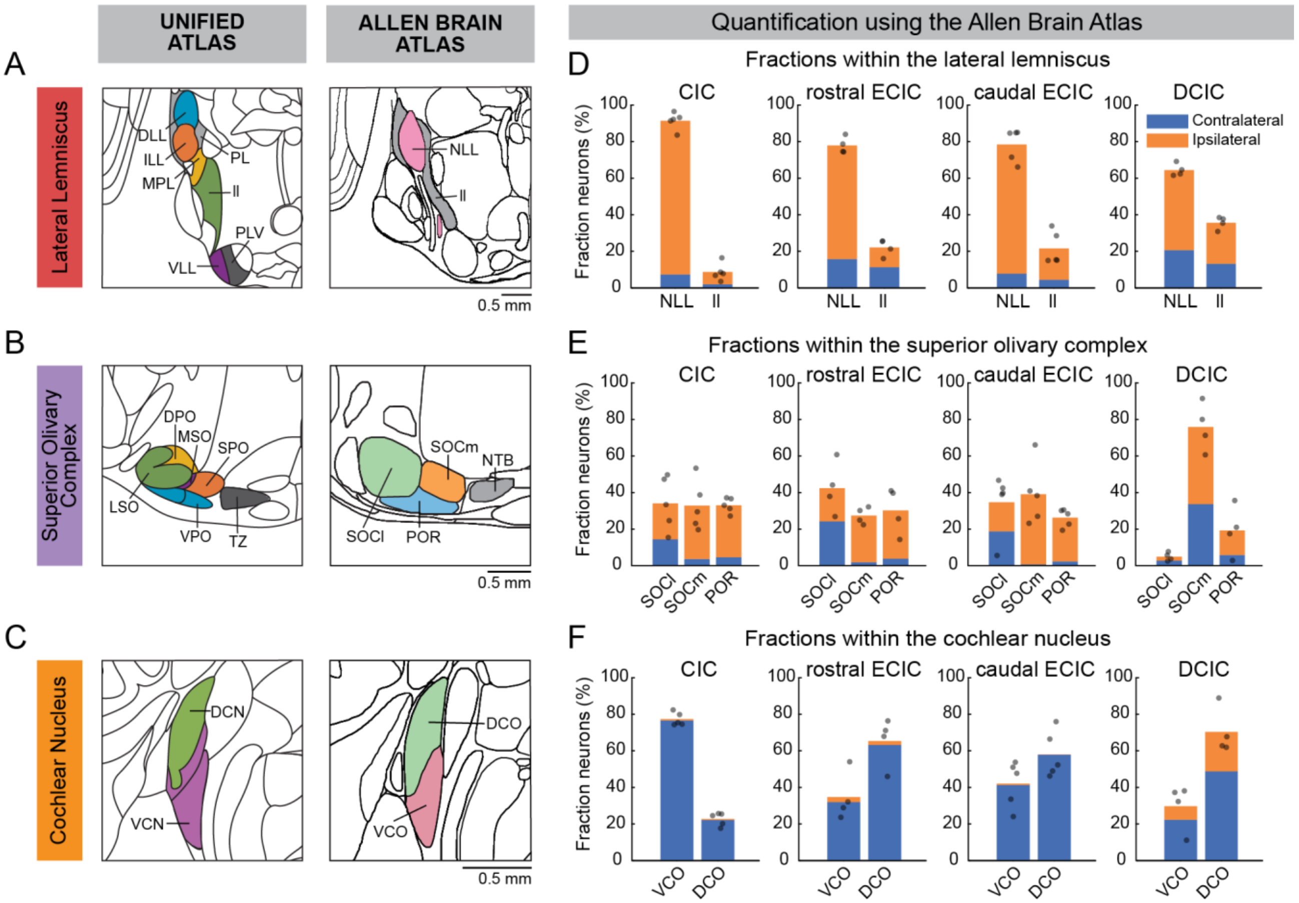


**Figure S1. Comparison between Kim's Unified Mouse Brain Atlas and the Allen Common Coordinate Framework.**

**A,** Schematic diagrams showing regional boundaries around the lateral lemniscus for the Unified Atlas (left) and Allen CCF (right). **B,** Same as in A, for the superior olivary complex. **C,** Same as in A, for the cochlear nucleus. **D,** Bar plots showing the fraction of presynaptic neurons in the NLL and ll (Allen CCF) for injections into the CIC (*n* = 5), rostral ECIC (*n* = 4), caudal ECIC (*n* = 5), and DCIC (*n* = 4). Stacked bars indicate ipsilateral (orange) and contralateral (blue) hemispheres. Individual dots represent individual animals. This plot corresponds to Fig. 4A generated using the Unified Atlas. DLL, ILL, VLL, and PL in the Unified Atlas are largely encompassed within the NLL in the Allen CCF. **E,** Same as in D, for presynaptic neurons in the SOCl, SOCm, and POR. This plot corresponds to Fig. 6A generated using the Unified Atlas. The coarse segmentation in the Allen CCF does not align well with histologically identified structures. **F,** Same as in D, for presynaptic neurons in the VCO and DCO. This plot corresponds to Fig. 8A generated using the Unified Atlas. AVCN and PVCN in the Unified Atlas are combined as VCO in the Allen CCF. In addition, the dorsal portions of AVCN and PVCN are included in the DCO of the Allen CCF.

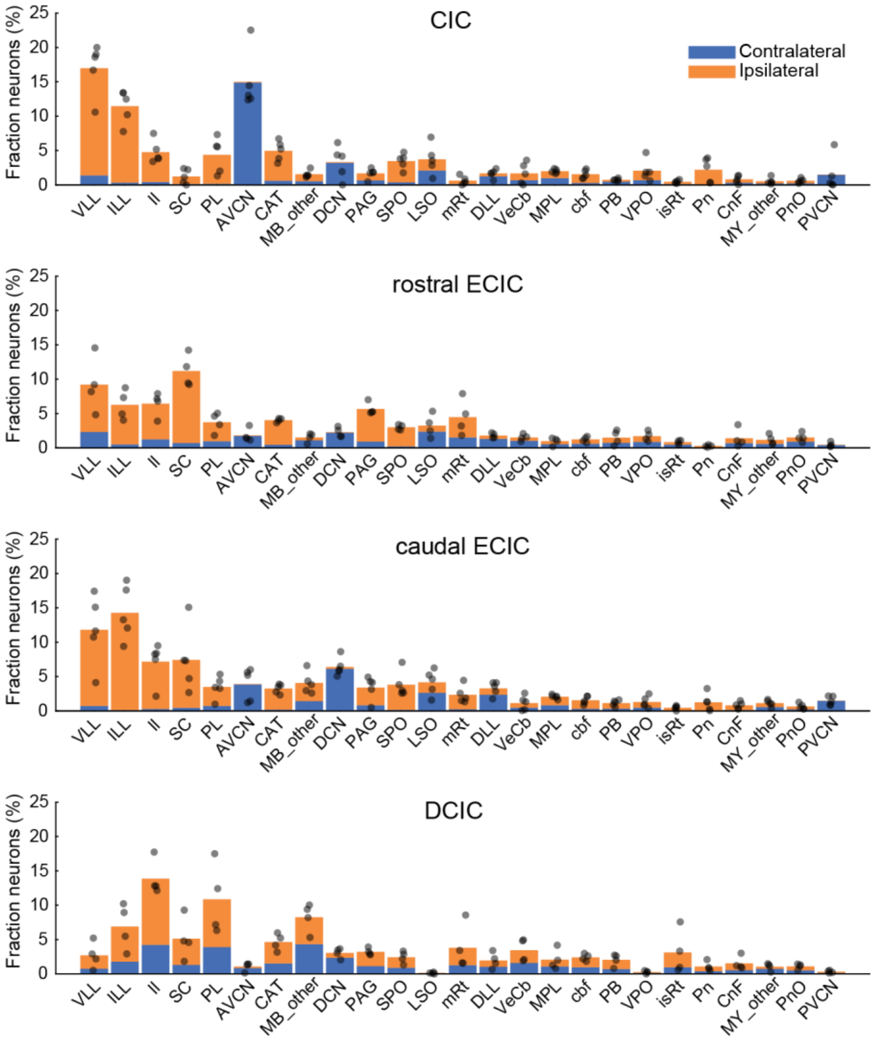


**Figure S2. Top 25 input regions across all IC injections.**

Bar plots showing the fraction of presynaptic neurons in the top 25 ascending input regions, ranked by total input fraction pooled across all four IC injection sites. Top, inputs to the CIC (n = 5). Second row, rostral ECIC (n = 4). Third row, caudal ECIC (n = 5). Bottom row, DCIC (n = 4). Stacked bars indicate ipsilateral (orange) and contralateral (blue) hemispheres. Dots represent individual animals.
Abbreviations: VLL, ventral nucleus of the lateral lemniscus; ILL, intermediate nucleus of the lateral lemniscus; ll, lateral lemniscus (fiber tract); SC, superior colliculus; PL, paralemniscal nucleus; AVCN, ventral cochlear nucleus, anterior part; CAT, nucleus of the central acoustic tract; MB_other, other midbrain regions; DCN, dorsal cochlear nucleus; PAG, periaqueductal gray; SPO, superior paraolivary nucleus; LSO, lateral superior olive; mRt, mesencephalic reticular formation; DLL, dorsal nucleus of the lateral lemniscus; VeCb, vestibulocerebellar nucleus; MPL, medial paralemniscal nucleus; cbf, cerebellum-related fiber tracts; PB, parabrachial nucleus; VPO, ventral periolivary nucleus; isRt, isthmic reticular formation; Pn, pontine nuclei; CnF, cuneiform nucleus; MY_other, other medullary regions; PnO, pontine reticular nucleus, oral part; PVCN, ventral cochlear nucleus, posterior part.

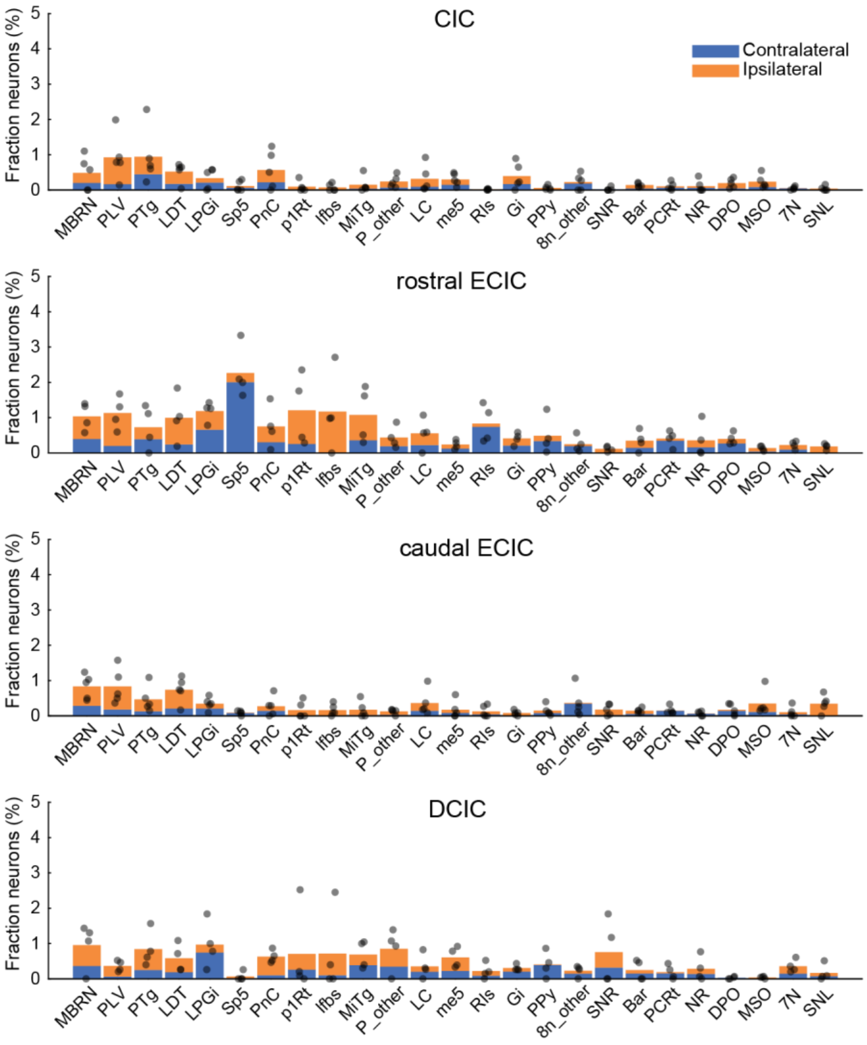


**Figure S3. Top 26-50 input regions across all IC injections.**

Bar plots showing the fraction of presynaptic neurons in the regions ranked 26–50 ascending input regions, ranked by total input fraction pooled across all four IC injection sites. Top, inputs to the CIC (n = 5). Second row, rostral ECIC (n = 4). Third row, caudal ECIC (n = 5). Bottom row, DCIC (n = 4). Stacked bars indicate ipsilateral (orange) and contralateral (blue) hemispheres. Dots represent individual animals.

Abbreviations: MBRN, midbrain raphe nuclei; PLV, paralemniscal nucleus, ventral part; PTg, pedunculotegmental nucleus; LDT, laterodorsal tegmental nucleus; LPGi, lateral paragigantocellular nucleus; Sp5, spinal trigeminal nucleus; PnC, pontine reticular nucleus, caudal part; p1Rt, prosomere 1 reticular formation; lfbs, lateral forebrain bundle system; MiTg, microcellular tegmental nucleus; P_other, other pontine regions; LC, locus coeruleus; me5, mesencephalic trigeminal tract; RIs, retroisthmic nucleus; Gi, gigantocellular reticular nucleus; PPy, parapyramidal nucleus; 8n_other, vestibulocochlear nerve (other components); SNR, substantia nigra, reticular part; Bar, Barrington nucleus; PCRt, parvicellular reticular nucleus; NR, nucleus raphe; DPO, dorsal periolivary region; MSO, medial superior olive; 7N, facial nucleus; SNL, substantia nigra, lateral part.

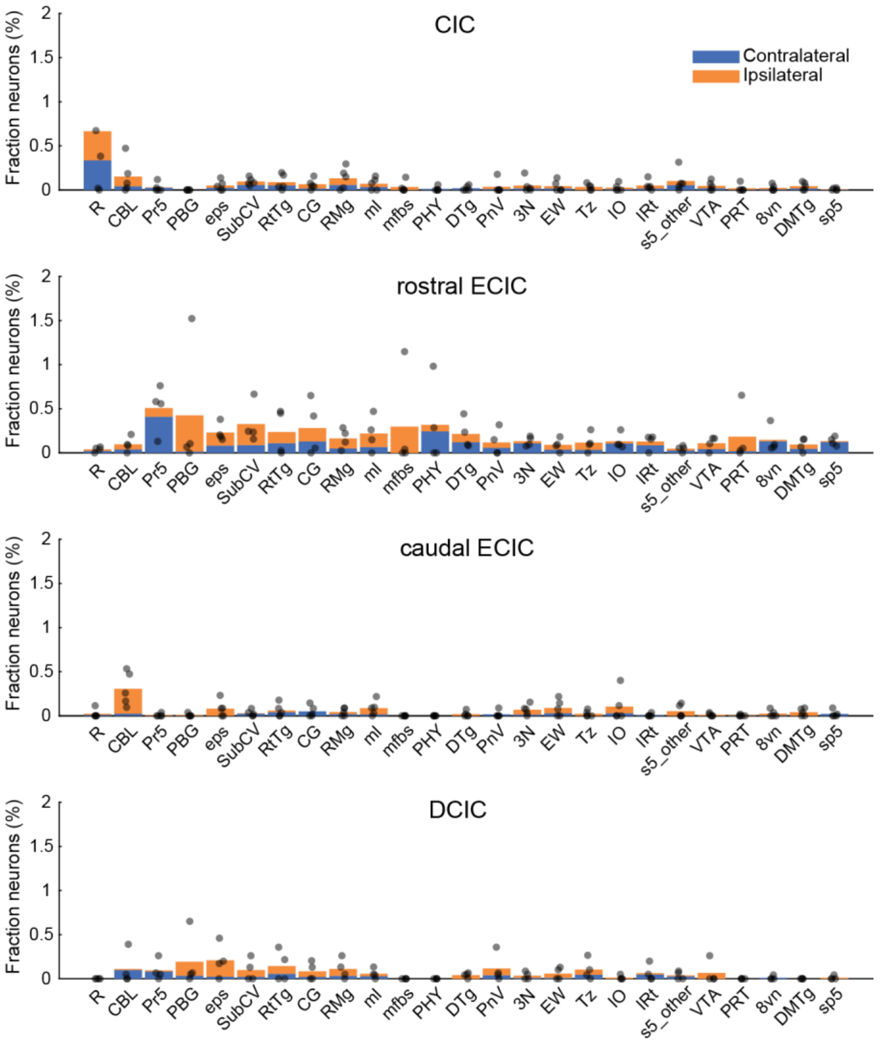


**Figure S4. Top 51-75 input regions across all IC injections.**

Bar plots showing the fraction of presynaptic neurons in the regions ranked 51–75 ascending input regions, ranked by total input fraction pooled across all four IC injection sites. Top, inputs to the CIC (n = 5). Second row, rostral ECIC (n = 4). Third row, caudal ECIC (n = 5). Bottom row, DCIC (n = 4). Stacked bars indicate ipsilateral (orange) and contralateral (blue) hemispheres. Dots represent individual animals.

Abbreviations: R, red nucleus; CBL, cerebellum; Pr5, principal sensory trigeminal nucleus; PBG, parabigeminal nucleus; eps, extrapyramidal fiber systems; SubCV, subcoeruleus nucleus, ventral part; RtTg, reticulotegmental nucleus of the pons; CG, central gray; RMg, raphe magnus nucleus; ml, medial lemniscus; mfbs, medial forebrain bundle system; PHY, perihypoglossal nuclei; DTg, dorsal tegmental nucleus; PnV, pontine reticular nucleus, ventral part; 3N, oculomotor nucleus; EW, Edinger–Westphal nucleus; Tz, nucleus of the trapezoid body; IO, inferior olivary nucleus; IRt, intermediate reticular nucleus; s5_other, sensory root of the trigeminal nerve, other parts; VTA, ventral tegmental area; PRT, pretectal region; 8vn, vestibular root of the vestibulocochlear nerve; DMTg, dorsomedial tegmental area; sp5, spinal trigeminal tract.

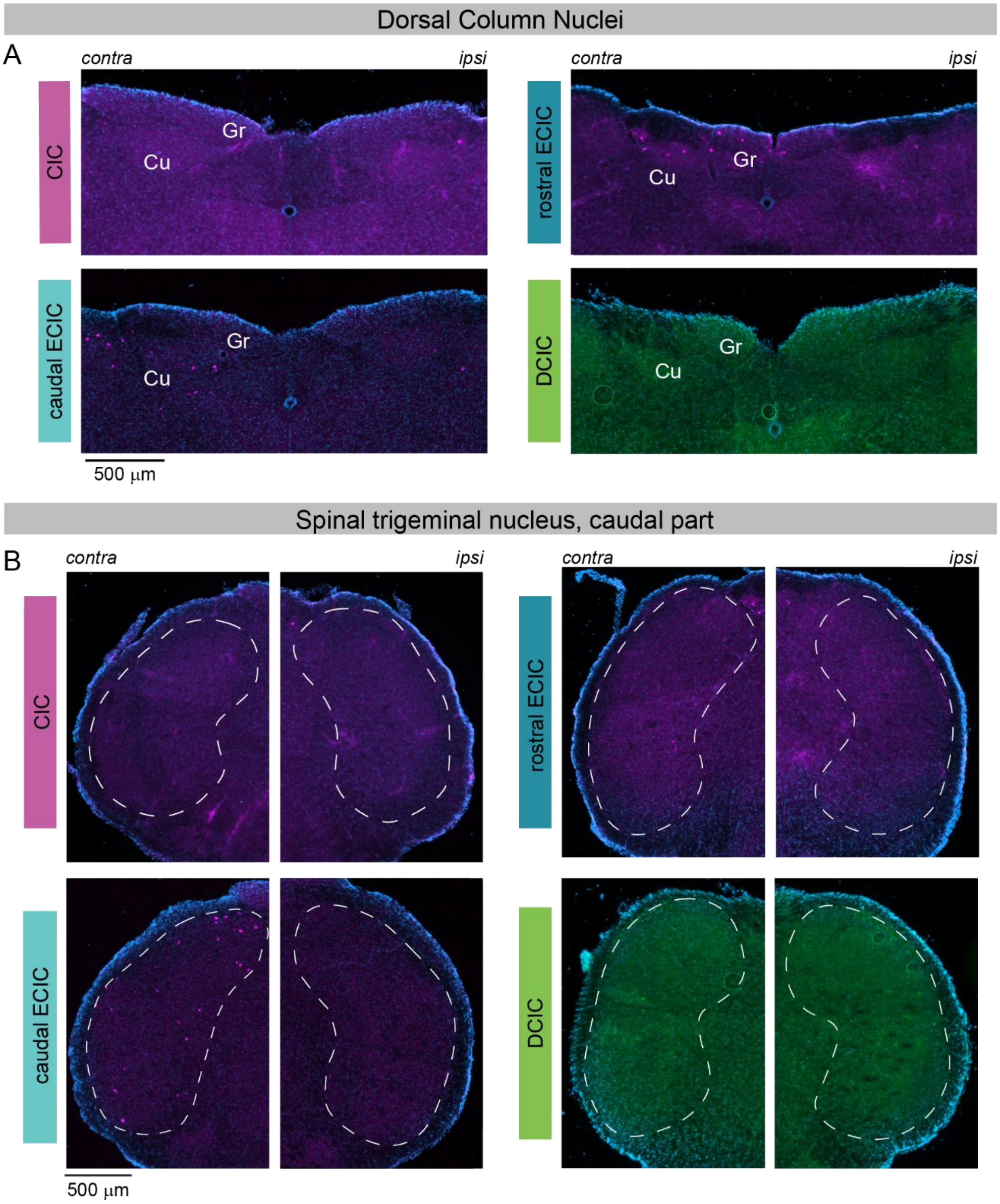


**Figure S5. Projections from the caudal medulla.**

**A,** Representative coronal sections showing labeling in the dorsal column nuclei following injections of WGA-647 (magenta) into the CIC (top left), rostral ECIC (top right), and caudal ECIC (bottom left), and WGA-488 (green) into the DCIC (bottom right). Gr, gracile nucleus; Cu, cuneate nucleus. Sections were counterstained with DAPI (blue). Labeling is observed in the dorsal column nuclei following injections into the rostral and caudal ECIC, but not the CIC or DCIC. **B,** Same as in A, for the caudal part of the spinal trigeminal nucleus (Sp5). Labeling is observed only following injection into the caudal ECIC.
